# A Beneficial Endornavirus Enhances the Fitness of Phytopathogenic Fungus *Rhizotonia solani*

**DOI:** 10.64898/2026.01.23.701230

**Authors:** Tianxing Pang, Bokang Li, Qianqian Sun, Zhiping Deng, Chunmei Cao, Zhensheng Kang, Ida Bagus Andika, Liying Sun

## Abstract

Although viruses are primarily characterized as pathogenic agents, certain viruses confer advantages to their hosts. The extent to which a virus can improve host biological performance, however, remains a fascinating topic in virology. An endornavirus, Rhizoctonia solani endornavirus IM (RsEV-IM), was identified as prevalent in *Rhizoctonia solani* isolates obtained from potato plants. Comparative analysis with a virus-free isogenic strain demonstrated that RsEV-IM infection enhances mycelial growth, sclerotium formation, stress tolerance, and fungal virulence across multiple plant species. Inoculation tests involving numerous *R. solani* strains confirmed that only RsEV-IM-infected strains exhibited high pathogenicity, independent of other mycovirus infections. Additionally, its secreted protein fraction of RsEV-IM-infected fungus contained elevated levels of various proteins, including those involved in cell wall degradation. This fraction not only facilitated *R. solani* infection but also suppressed the growth of other fungi and bacteria. These findings position RsEV-IM as a beneficial virus that widely enhances its host’s biological fitness. From both pathological and ecological perspectives, these observations are significant, as they reveal that a mycovirus can serve as a key virulence determinant in fungal populations and potentially shape microbial community dynamics in natural environments.

**Importance:** Fungal pathogenicity and ecological traits have long been thought to be primarily governed by endogenous genetic factors. However, this study reveals a mutualistic relationship between an endornavirus (RsEV-IM) and *Rhizoctonia solani*, demonstrating that viral infection enhances fungal virulence and ecological fitness. RsEV-IM stimulates fungal growth and the secretion of cell wall-degrading enzymes, resulting in a severe disease phenotype. Ecologically, RsEV-IM-infected fungi potentially gain a competitive advantage over soil microbiota. These findings present a key example of a virus acting as an essential extrachromosomal determinant of fungal pathogenicity and ecosystem interactions. Our results advance the understanding of fungal virulence mechanisms and underscore the broader significance of beneficial virus-fungus associations in agriculture and microbial ecology.

## Introduction

Pathogenic interactions between viruses and their hosts are often more emphasized due to the detrimental impacts of viruses on human and animal health, as well as agricultural production (1, 2). In fact, extensive studies of viruses infecting various organisms have revealed that cryptic and/or persistent infections are more common, often suggesting a commensalistic relationship with their hosts (3–5). On the other hand, a growing number of studies have demonstrated mutualistic virus-host interactions, in which viruses provide certain benefits to their hosts or directly promote advantageous host phenotypes (6). Nevertheless, the impacts of the majority of viruses on their hosts remain largely obscure, highlighting the need for further extensive investigation to deepen our understanding of the cellular and ecological roles of viruses in nature.

A vast number of mycoviruses have been identified from diverse fungal taxa (7). Although the majority of mycoviruses appear to be asymptomatic, many have been observed to negatively affect host phenotypes. In particular, numerous mycoviruses identified in phytopathogenic fungi attenuate fungal virulence (8). Notably, a number of mycoviruses have also been found to be beneficial to their hosts (9). The enhancement of fungal virulence phenotypes (hypervirulence) by mycovirus infection has been documented in some plant, insect and mammalian pathogenic fungi (10–16). However, it remains unclear whether these mycovirus-induced hypervirulence effects are specific to fungal strains with particular genetic backgrounds or whether a mycovirus could serve as a key virulence determinant across fungal populations in nature.

Endornaviruses are members of the family *Endornaviridae*, characterized by non-encapsidated, linear, positive-sense RNA genomes containing a single open reading frame (ORF) that encodes a large polyprotein (17). Endornaviruses were initially identified in crop plants such as bean, rice, and pepper, with general characteristics of persistent and cryptic infections (18). Subsequently, endorna or endorna-like viruses have also been identified in fungi and oomycetes (19). Currently, the family *Endornaviridae* is divided into two officially recognized genera, *Alphaendornavirus* and *Betaendornavirus*, based on genome size and phylogenetic relationships (17) while a novel genus, Gammaendornavirus, was recently proposed (20). The effects of endornavirus infection on fungal hosts have not been clearly demonstrated, although some studies have suggested antagonistic effects on growth, stress tolerance, and virulence (21, 22). A recent study showed an enhancing effect of co-infection with two endornaviruses on the virulence of the oomycete host (23).

*Rhizoctonia solani* Kühn (teleomorph: *Thanatephorus cucumeris*) is a necrotrophic, soil-borne phytopathogenic fungus that affects a wide range of economically important crop plants worldwide (24). *R. solani* is a fungal species complex whose subspecific groups are classified according to hyphal anastomosis interactions, termed anastomosis groups (AGs) (25). In potato, *R. solani* causes stem canker or black scurf diseases (26). Among the several AGs known to infect potato, AG-3PT (potato type), a subgroup of AG-3, is considered the predominant and most aggressive causal agent of Rhizoctonia disease in potatoes (27). A number of candidate genes potentially involved in *R. solani* pathogenicity have been identified (28, 29). However, the difficulty in transforming *R. solani* has impeded further detailed molecular and genetic studies on its pathogenicity. Diverse mycoviruses have been identified in *R. solani*, but their impacts on the host remain largely unknown (30).

In this study, we examined the effects of infection by an endornavirus named *Rhizoctonia solani* endornavirus IM (RsEV-IM) in *R. solani.* RsEV-IM infection was observed to promote fungal growth and appeared to be a determining factor for fungal virulence. Our results demonstrate a mutualistic interaction between a mycovirus and a phytopathogenic fungus, with significant pathological and potential ecological impacts.

## Results

### An endornavirus (RsEV-IM) with growth-promoting effects is prevalent in *R. solani* strains

We collected a large number of *R. solani* strains isolated from potato plants grown in different areas of Inner Mongolia Autonomous Region, China (Fig. S1). Total RNAs extracted from 101 fungal strains were pooled and subjected to next-generation sequencing (NGS) and subsequent bioinformatic analyses. Numerous virus-related sequences were obtained from the dataset. One large virus contig with a complete open reading frame (ORF, contig DN1616, 18589 nt) was particularly notable due to its high read numbers (Fig. S2). BLASTX analysis revealed that this contig had sequence similarity to endornaviruses, with the highest identity (24.34%) to *Rhizoctonia solani* Endornavirus 5 (Acc. No. QDW65432.1). The complete viral genome sequence was obtained through rapid amplification of cDNA ends (RACE) and RT-PCR and deposited in GenBank (Acc. No. OP763640). This virus is tentatively named *Rhizoctonia solani* Endornavirus IM (RsEV-IM).

The RsEV-IM genome is 18,636 nt in length, with a nick identified at the nt position 89 in the coding region through 5’ RACE (Fig. 1A). Many members of the genus *Alphaendornavirus* contain a site-specific nick near the 5′ end of the genome, although its implication in virus replication remains unknown (18). The RsEV-IM genome consists of 5’ and 3’ untranslated regions (61 nt and 47 nt, respectively) flanking a large ORF encoding a potential polyprotein with conserved methyltransferase (MTR), helicase (HEL), and RNA-dependent RNA polymerase (RdRp) motifs (Fig. 1A). Additionally, a putative cysteine-rich region (CRR) with three conserved signature sequences “CXCC” (20), was identified between the MTR and HEL domains (Fig. S3). Phylogenetic analysis based on the RdRp domain sequences of selected endornaviruses and endorna-like viruses showed that RsEV-IM clusters with members of the genus *Alphaendornavirus* (Fig. S4). Multiple sequence alignment revealed that the RdRp domain of RsEV-IM shares six conserved motifs with those of alphaendornaviruses (Fig. S5). Notably, although RsEV-IM exhibits the most similarity in RdRp domain sequences and genome size (>11.9 kb) to those of alphaendornaviruses, it encodes an MTR domain typically found in betaendornaviruses (18). Overall, our analyses suggest that RsEV-IM is a candidate member of the genus *Alphaendornavirus*.

**Figure 1.**
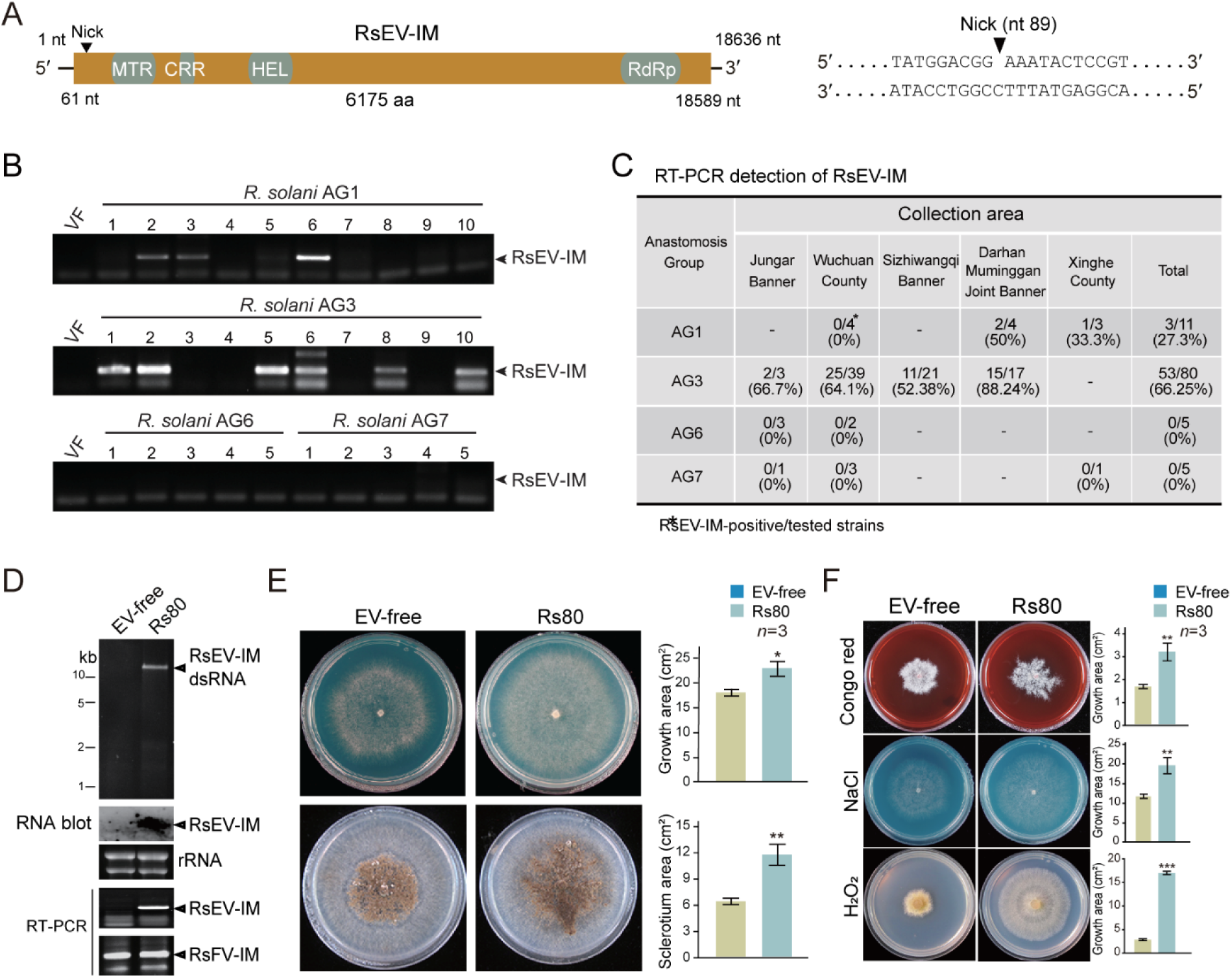
Molecular and biological characteristics of RsEV-IM. **(A)** Schematic representation (not to scale) of genome structure of RsEV-IM. Colored boxes represent open reading frames (ORFs) and black lines represent 5’- and 3’-untranslated regions (UTRs). Nucleotide positions of ORFs and UTRs are indicated. The relative positions of conserved domains in the encoded protein are shown in ORF (colored capsule-shaped forms): RNA dependent RNA polymerase (RdRp), helicase (Hel), cysteine-rich region (CRR), and methyltransferase (MTR). The sequences surrounding a nick at the nt position 89 in the coding region were presented. **(B)** The representative results of RT-PCR detection of RsEV-IM in different *R. solani* AG strains. **(C)** Summary of RsEV-IM prevalence in *R. solani* strains obtained from different areas in Inner Mongolia as determined by RT-PCR detection. **(D)** Detection of virus in Rs80 and RsEV-IM-cured strains (EV-free) by dsRNA, RNA blot, and RT-PCR analyses. **(E)** Phenotypic growth and sclerotium formation of Rs80 and EV-free strains on PDA medium (60 mm plate; imaged at 3 days). Colony and sclerotium areas (mean ± SD, *n*=3) are shown. Asterisks indicate significance (\**P* < 0.05, \*\**P* < 0.01, \*\*\**P* < 0.001; Student’s t-test). **(F)** Phenotypic growth of Rs80 and EV-free strains on PDA medium (60 mm plate; imaged at 3 days) supplemented with stress-inducing agents (congo red; cell wall stress, NaCl; osmotic stress, H_2_O_2_; oxidative stress). Colony and sclerotium areas (mean ± SD, *n*=3) are shown (asterisks as above).

Analysis of the internal transcribed spacer (ITS) region indicated that the isolated fungal strains (101 strains) predominantly belong to the AG-3 group (79%), while the remaining strains belong to the AG-1, AG-6, and AG-7 groups. RT-PCR detection was performed to investigate the prevalence of RsEV-IM in the fungal strains. The results revealed that 66% of AG-3 strains and 27% of AG-1 strains were positive for RsEV-IM infection, whereas none of the AG-6 and AG-7 strains tested positive for RsEV-IM (Fig. 1B and C). It was also noted that RsEV-IM-infected AG-3 strains were obtained from every area where fungal strains were collected (Fig. 1C), suggesting that RsEV-IM is highly prevalent and widely distributed in potato-growing areas of Inner Mongolia region.

To further study RsEV-IM in detail, one fungal strain (Rs80, AG-3) was selected due to its single dsRNA profile (Fig. S6). However, further NGS analysis revealed that the Rs80 strain was also infected with a novel fusarivirus (a positive-sense RNA virus, family *Fusariviridae*), tentatively named *Rhizoctonia solani* fusarivirus IM (RsFV-IM, GenBank Acc. No. PV941035, Fig. S7). In attempts to cure Rs80 of RsEV-IM infection, we performed multiple rounds of hyphal tipping (Fig. S8), one RsEV-IM-free isogenic strain (referred to as EV-free) was obtained, as confirmed by dsRNA, RNA blot, and RT-PCR analyses, although the strain still retained RsFV-IM infection (Fig. 1D). Interestingly, the EV-free strain exhibited a significantly slower growth rate and reduced sclerotium formation compared to the Rs80 strain when cultured on PDA medium (Fig. 1E). RsEV-IM-infected fungal strains regenerated from sclerotia (Fig. S9), suggesting that RsEV-IM can be stably maintained during long-term survival of *R. solani* under harsh environmental conditions. Moreover, the EV-free strain also showed smaller colonies than the Rs80 strain when cultured on PDA medium supplemented with stress-inducing agents or fungicides (Fig. 1F and Fig. S10). These observations suggest that RsEV-IM infection promotes the growth and stress tolerance of the fungal host *R. solani*.

### RsEV-IM infection is associated with the strong pathogenicity of *R. solani* strains

To examine the effect of RsEV-IM infection on *R. solani* pathogenicity in potato and other plant species, Rs80 and EV-free strains were inoculated into the lower stems of young potato, tomato, and *N. benthamiana* plants, as well as young seedlings of pepper, cucumber, watermelon, and radish. Strikingly, at 7 days post-inoculation (dpi), the Rs80 strain induced severe stem rot in all tested plants, causing heavy rotting of the stems and eventual plant death. In contrast, plants inoculated with the EV-free strain remained vigorous, developing only small fungal lesions at the inoculation sites (Fig. 2A). In a separate experiment, potato tuber seeds were planted in soil artificially infested with Rs80 or EV-free strain mycelia. By 14 days after planting, all potato plants grown in Rs80-infested soil had died, whereas those in EV-free-infested soil remained asymptomatic and healthy (Fig. 2B). These results demonstrate that RsEV-IM infection significantly enhances the virulence of *R. solani* strain Rs80.

**Figure 2.**
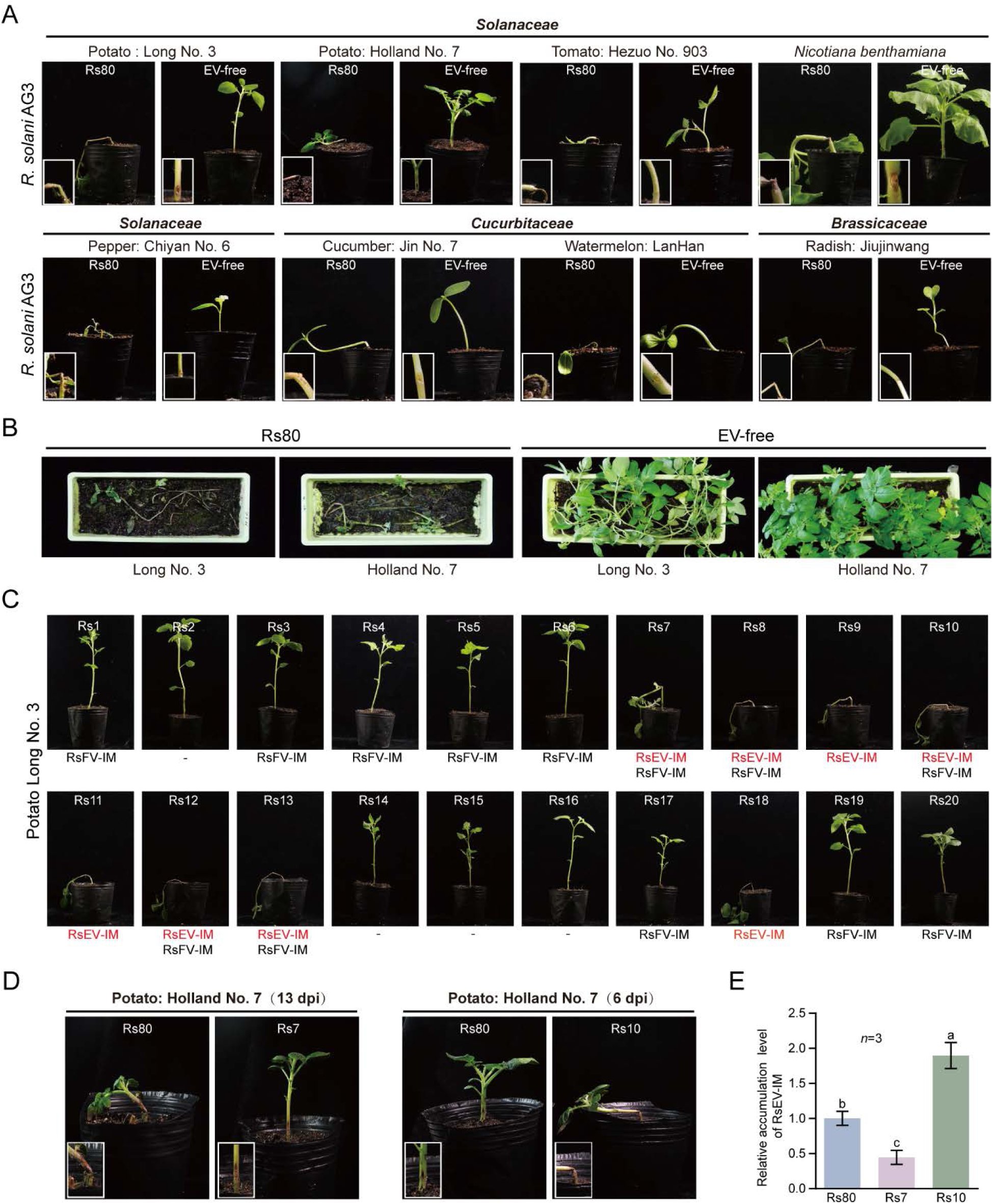
Effects of RsEV-IM infection on pathogenicity of *R. solani* strains. **(A)** Development of stem rot in lower stems of potato and other plant species inoculated with Rs80 and EV-free strains. Insets present close-up views of the inoculated stems (Magnification: 5×). Plants were photographed at 7 days post-inoculation (dpi). The presented image is representative of three inoculated plants. **(B)** Potato seeds grown on soils artificially infested with Rs80 and EV-free strains. Plants were photographed at 14 days after planting. **(C)** Development of stem rot in lower stems of potato plants inoculated with different *R. solani* AG-3 strains. Plants were photographed at 7 dpi. The presence of RsEV-IM and/or RsFV-IM in each fungal strain is indicated below the plant image. **(D)** Development of stem rot in lower stems of potato plants inoculated with Rs80, Rs7, and Rs10 strains. The presented image is representative of three inoculated plants. **(E)** Relative RsEV-IM accumulation levels in Rs80, Rs7, and Rs10 strains analyzed by qRT-PCR (mean ± SD, *n*=3). The value from the Rs80 sample was normalized to 1.00. Different letters indicate significance (*P* < 0.05, one-way ANOVA).

To further investigate whether the association between RsEV-IM infection and strong pathogenicity is widespread among *R. solani* strains, we conducted fungal inoculation assays using 40 *R. solani* strains (all AG-3) collected in this study. These strains appear to harbor different mycoviruses as indicated by heterogenous dsRNA profiles (Fig. S6). The presence or absence of RsEV-IM and RsFV-IM in the fungal strains was confirmed by RT-PCR. These strains were inoculated onto the seven plant species described earlier. Overall, the inoculation tests revealed a strong correlation between RsEV-IM infection and the induction of severe stem rot (plant death) across plant species, irrespective of RsFV-IM presence (Fig. 2C and Table S1). However, one discrepancy was observed: strain Rs7, despite harboring RsEV-IM, did not consistently induce severe stem rot in all tested plant species (Fig. 2D and Table S1). Additionally, strain Rs10 stood out due to its faster induction of plant death compared to other strains (Fig. 2D). Reverse transcription quantitative polymerase chain reaction (RT-qPCR) analysis showed that RsEV-IM accumulation was lower in Rs7 than in Rs80, whereas it was higher in Rs10 than in Rs80 (Fig. 2E). This suggests that RsEV-IM accumulation levels influence the virulence potential of *R. solani*. In addition, fungal inoculation assays using both RsEV-IM-free and -infected AG-1 strains also showed correlation between RsEV-IM infection and severe stem rot (Fig. S11). Together, these findings suggest that RsEV-IM plays a determining role in *R. solani* pathogenicity in wide variety of plant species.

### Virus inoculation and RNA interference (RNAi) confirms that RsEV-IM is responsible for enhanced growth and pathogenicity of *R. solani* strains

To further confirm that RsEV-IM is the causative agent for the enhanced biological traits in *R. solani*, we attempted to inoculate EV-free and other RsEV-IM-free fungal strains (Rs2, Rs3 and Rs4) by directly applying total RNA extracted from the Rs80 strain to the fungal mycelia (Fig. 3A). After successive subculturing of the treated mycelia, RsEV-IM was stably detected in these fungal strains via RT-PCR and dsRNA extraction (Fig. 3B), indicating successful viral inoculation through this method. Consistent with our previous observations, the RsEV-IM-infected strains exhibited enhanced growth, high virulence, and increased stress tolerance (Fig. 3C, 3D, and Fig. S12).

**Figure 3.**
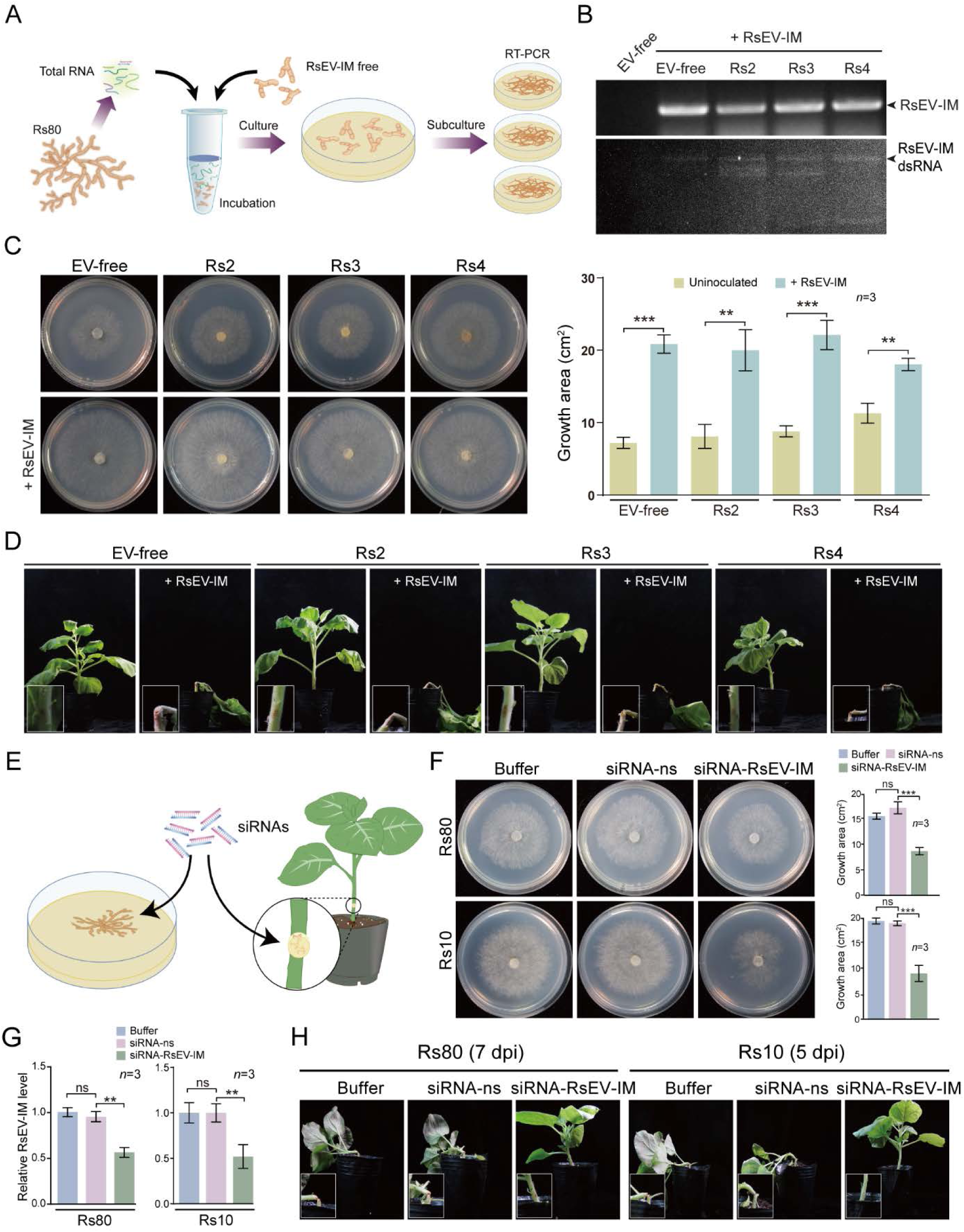
Inoculation of RsEV-IM and RNAi-mediated suppression of RsEV-IM. **(A)** An illustration depicting the experimental procedure for inoculation of fungal strains with RsEV-IM. **(B)** Detection of RsEV-IM in fungal strains after virus inoculation by RT-PCR and dsRNA analyses. **(C)** Mycelial growth of fungal strains that had been inoculated with RsEV-IM on PDA medium (60 mm plate; imaged at 3 days). Colony areas (mean ± SD, *n*=3) are shown. Asterisks indicate significance (\*\**P* < 0.01, \*\*\**P* < 0.001; Student’s t-test). **(D)** Development of stem rot in lower stems of *N. benthamiana* inoculated with fungal strains that had been inoculated with RsEV-IM. Insets present close-up views of the inoculated stems (Magnification: 5×). Plants were photographed at 10 days post-inoculation (dpi). The presented image is representative of three inoculated plants. **(E)** An illustration of the procedure for applying exogenous siRNAs to fungal mycelia, either cultured on PDA medium or inoculated on a plant’s lower stem. **(F)** Mycelial growth of Rs80 and Rs10 strains treated with nonspecific (ns) or RsEV-IM-specific siRNAs on PDA medium (60 mm plate; imaged at 3 days). Colony areas (mean ± SD, *n*=3) are shown (asterisks as above). **(G)** Relative RsEV-IM accumulation levels in Rs80 and Rs10 strains treated with non-specific (ns) or RsEV-IM-specific siRNAs analyzed by qRT-PCR (mean ± SD, *n*=3, asterisks as above). The value from buffer treated sample was normalized to 1.00. **(H)** Development of stem rot in lower stems of *N. benthamiana* plants inoculated with Rs80 and Rs10 strains treated with ns or RsEV-IM-specific siRNAs. Insets present close-up views of the inoculated stems (Magnification: 5×). Plants were photographed at 5 and 7 dpi. The presented image is representative of three inoculated plants.

Next, we further examined whether RsEV-IM accumulation levels are correlated with the virulence potential of *R. solani* (Fig. 2E). The application of exogenous siRNAs or dsRNA has been successfully used to induce RNAi-mediated suppression against endogenous fungal genes (31–33). To examine whether this approach could be applied to reduce RsEV-IM accumulation in *R. solani*, we treated the Rs80 and Rs10 strains with synthetic siRNAs complementary to the RsEV-IM genome (Fig. 3E). Treatment with RsEV-specific siRNAs, but not nonspecific (ns) siRNAs, markedly reduced Rs80 and Rs10 growths (Fig. 3F) and was associated with reduced RsEV-IM accumulation (Fig. 3G). Furthermore, applying RsEV-specific siRNAs to the Rs80 and Rs10 strains inoculated into *N. benthamiana* stems diminished stem rot induction (Fig. 3H). These results further confirm the positive role of RsEV-IM in *R. solani* growth and pathogenicity, and also open an alternative means to control *R. solani*-induced crop diseases by targeting RsEV-IM.

### The secreted protein fraction from RsEV-IM-infected fungus promotes stem rot induction and exhibits antimicrobial activities

Fungi secrete diverse groups of proteins that facilitate fungal growth and host colonialization (34). To investigate the role of secreted proteins in the enhanced pathogenicity of *R. solani* following RsEV-IM infection, we isolated secreted proteins from the Rs80 and EV-free strains and applied them to potato shoots alongside inoculation with the EV-free strain. The application of secreted protein fraction from Rs80, but not from the EV-free strain, induced severe stem rot (Fig. 4A). Notably, the Rs80 secreted proteins alone, even in the absence of fungal inoculation, triggered stem rot-like symptoms, whereas the EV-free fraction did not (Fig. 4B). These results indicate that specific components within the Rs80 secreted protein fraction contribute to stem rot induction.

**Figure 4.**
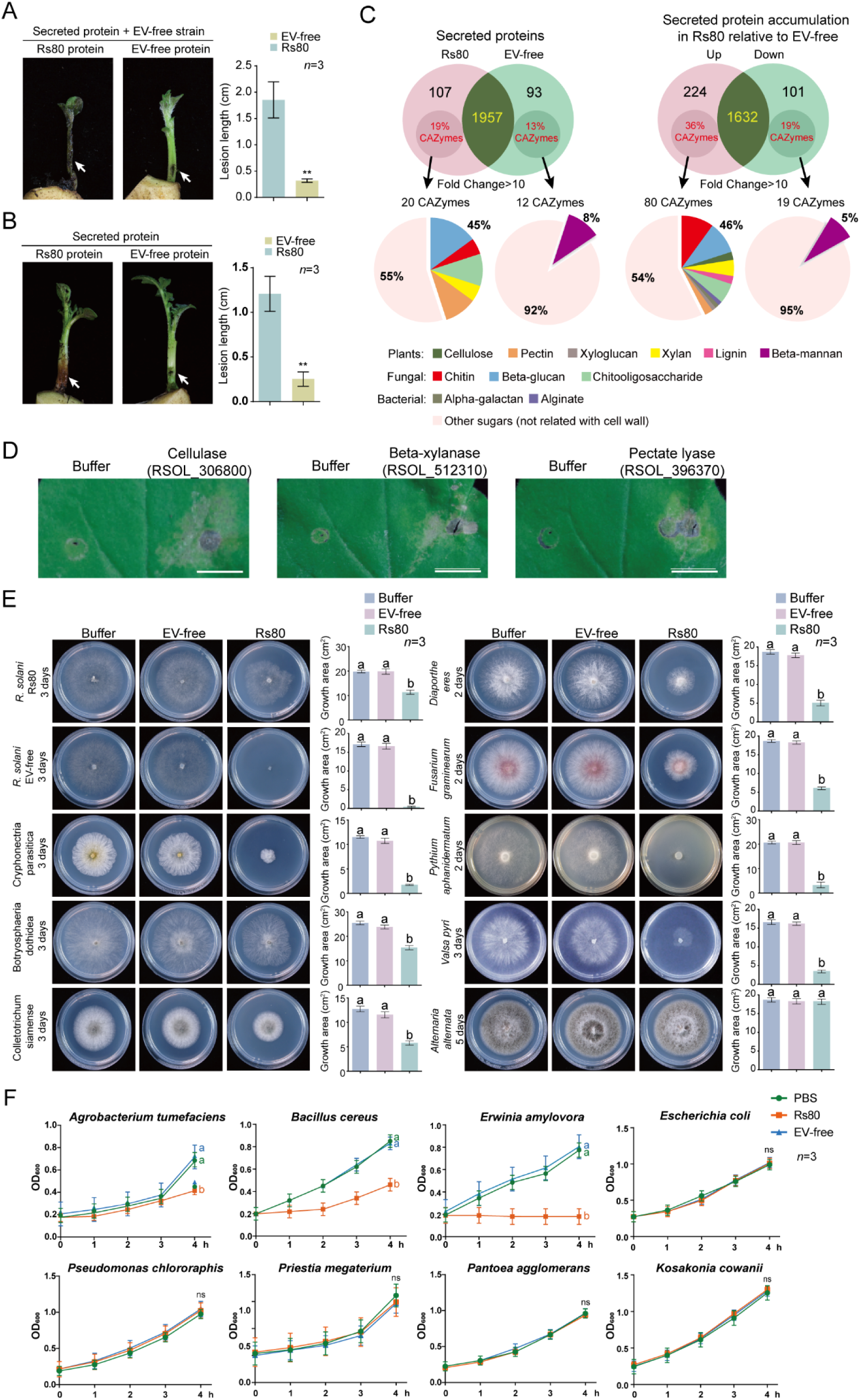
Effects and compositions of secreted protein fraction from *R. solani* strains. **(A)** Development of stem rot in potato shoots inoculated with EV-free strain with addition of secreted protein fraction from Rs80 or EV-free strain (upper panel) or in potato shoots treated with secreted protein fraction from either strain (lower panel). Arrows indicate the position of fungal inoculation and/or secreted protein treatment. Shoots were photographed at 7 dpi. The presented image is representative of three inoculated shoots. Lesion length measured from the shoots (mean ± SD, *n*=3) are shown. Asterisks indicate significance (\**P* < 0.05, \*\**P* < 0.01, \*\*\**P* < 0.001; Student’s t-test). **(B)** Compositions of secreted protein fractions from Rs80 or EV-free strains analyzed by LC-MS/MS. The proportions of secreted CWDEs with different abundances, targeting diverse substrates, were presented. **(C)** Leaf tissue treated with eukaryotically expressed *R. solani* CWDEs upregulated by RsEV-IM infection. Leaves were photographed at 3 days after treatment. **(D)** Mycelial growth of different fungal species treated with secreted protein fraction from Rs80 or EV-free strain on PDA medium (60 mm plate; imaged at 2−3 days). Colony areas (mean ± SD, *n*=3) are shown. Different letters indicate significant differences (*P* < 0.05, one-way ANOVA). **(E)** Growth rate of different bacterial species treated with secreted protein fraction from Rs80 or EV-free strain cultured on liquid medium. Data represent mean ± SD (*n*=3). The data at 4 h were subjected to statistical analysis (letters as above).

The Rs80 and EV-free secreted protein fractions were then analyzed by liquid chromatography-tandem mass spectrometry (LC-MS/MS) to investigate their protein compositions. A total of 2,175 proteins were identified, with 107 and 93 proteins exclusively expressed in the Rs80 and EV-free strains, respectively. Among the 1,957 secreted proteins common to both strains, 224 proteins showed a >10-fold increase in the Rs80 strain, while 101 proteins were reduced by >10-fold compared to the EV-free strain (Fig. 4C and Supplementary File S1).

Fungal basal secretomes contain various enzymes involved in the synthesis and breakdown of polysaccharides, commonly referred to as carbohydrate-active enzymes (CAZymes). Many of these act as cell wall-degrading enzymes (CWDEs) due to their ability to hydrolyze structural polysaccharide components of cell walls (35). Studies have demonstrated that CWDEs play a crucial role in the pathogenicity of diverse plant-pathogenic fungi (36–38). Notably, the genomes of necrotrophic or hemibiotrophic fungi encode more CAZymes than those of biotrophic fungi (39).

Using the dbCAN3 pipeline (40), we identified 20 and 12 CAZymes exclusively expressed in the Rs80 and EV-free strains, respectively. Strikingly, 9 (45%) of the Rs80-specific CAZymes were CWDEs, compared to only 1 (8%) in the EV-free strain. Among secreted proteins common to both strains at a >10-fold change threshold, we identified 80 CAZymes with increased accumulation and 19 with reduced accumulation in Rs80 relative to the EV-free strain. The increased accumulation group contained a substantially higher proportion of CWDEs than the reduced accumulation group (46% vs. 5%) (Fig. 4C and Supplementary File S2). This pattern of preferential enrichment of CWDEs in the increased accumulation group was consistently observed at both >5-fold and >1.5-fold change thresholds (Fig. S13). Notably, in Rs80 strain CWDEs with increased accumulation targeted diverse substrates, including plant cell wall components, fungal cell wall constituents and bacterial cell wall elements (Fig. 4C). Other proteins commonly found in fungal secretomes, such as proteases, oxidoreductases, and lipases, were present in the increased accumulation group (Supplementary File S2).

To verify the cell wall-digesting activities of these enzymes, three genes encoding cellulase, β-xylanase, and pectate lyase (showing 9.8-, 76.9-, and 205.5-fold increases in expression, respectively) were cloned and expressed in a eukaryotic system. The purified proteins were infiltrated into *N. benthamiana* leaves. Three days post-infiltration, leaf tissues treated with these proteins exhibited cell lysis and disruption (Fig. 4D), confirming cell wall degradation activity.

Since the secreted CWDEs also target fungal and bacterial cell wall components (Fig. 4C), we then tested the effect of secreted protein fractions on the growth of various fungal and bacterial species. Treatment with the secreted protein fraction from the Rs80 strain (but not that from the EV-free strain) severely inhibited the growth of the EV-free strain on PDA, while the same treatment also reduced the growth of the Rs80 strain but to a much lesser degree than that observed for the EV-free strain (Fig. 4E). These results suggest that RsEV-IM infection induces fungal tolerance against the high accumulation of CWDEs. Treatment of seven other different plant pathogenic fungi showed that Rs80 secreted proteins reduced growth in all species except for one *Alternaria* species (Fig. 4E). We next tested the effect of these secreted proteins on bacteria grown in liquid culture. Of the eight bacterial species tested, growth was inhibited in *Agrobacterium tumefaciens*, *Bacillus cereus*, and *Erwinia amylovora*, but not in the other five species (Fig. 4F). This indicates that the antibacterial spectrum of the Rs80 secreted proteins is relatively narrow compared to their antifungal spectrum. These observations indicate that RsEV-IM infection promotes the secretion of proteins with antimicrobial effects, particularly those with cell wall-degrading activities.

## Discussion

Inner Mongolia is one of China’s largest potato-producing regions. In recent years, it has been affected by high incidence rates of black scurf disease (41). AG-3 is the dominant group infecting potato plants in Inner Mongolia region and is thus the main anastomosis group causing Rhizoctonia disease in this area (41). In this study, *R. solani* strains were collected from potato plants across five planting areas in west and central Inner Mongolia. The majority of strains (79%) belonged to AG-3, and RsEV-IM was highly prevalent (52–88%) in AG-3 strains isolated from different regions. These results suggest a potential link between the high incidence and severity of stem rot or black scurf disease in Inner Mongolia and RsEV-IM infection in *R. solani*.

The extent of RsEV-IM distribution in *R. solani* populations, particularly those infecting potatoes, remains unclear. While numerous endornaviruses have been identified in *R. solani* strains infecting non-potato hosts (e.g., rice, tobacco), none show close genomic similarity to RsEV-IM (30, 42, 43). Metatranscriptomic analysis of binucleate and multinucleate *R. solani* strains infecting potatoes across various Chinese provinces/regions yielded multiple partial endornavirus-related sequence contigs, though these data are not publicly available (44). Notably, characterization of viruses in two highly and mildly virulent *R. solani* AG-3 strains isolated from potatoes in New Zealand, using random amplification of total dsRNA, obtained partial sequences with high homology (99%) to RsEV-IM (Fig. S14) (45). This suggests the possibility that RsEV-IM exists in potato-infecting AG-3 populations worldwide. Further extensive screening of RsEV-IM infection in *R. solani* strains from potato-growing regions is necessary to clarify the relationship between RsEV-IM infection and the incidence/severity of Rhizoctonia diseases.

Fungal pathogenicity and its general traits are traditionally thought to be determined by endogenous genetic factors (46). However, the discovery that fungi are commonly infected with diverse mycoviruses, many of which alter the phenotypes of their fungal hosts, challenges this view (47). Transcriptomic analyses of rice-infecting *R. solani* AG1-IA strains have identified potential virulence gene candidates, including those encoding putative secreted effectors, cell wall-degrading enzymes, proteases, and proteins involved in secondary metabolite production (28, 29, 48, 49). In contrast, virulence genes in *R. solani* AG-3 and other AGs remain understudied. Our observations show that RsEV-IM infection is associated with enhanced secretion of CWDEs as well as other proteins that potentially enhance fungal pathogenicity. This suggests that RsEV-encoded protein(s) do not directly act as virulence factors. Instead, RsEV-IM replication or its viral proteins appear to alter *R. solani*’s physiology and metabolism including through the reprogramming of gene expression. How RsEV-IM induces these changes, thereby improving the biological performance of *R. solani*, remains an intriguing topic for future molecular studies.

CWDEs have been explored for their antimicrobial applications (50, 51). Interestingly, RsEV infection leads to increased secretion of various CWDEs targeting cell wall components of plants, fungi, and bacteria, indicating a global upregulation or secretion of proteins involved in host colonization and competition against diverse microbes. The finding of increased CWDE accumulation in *R. solani* mediated by RsEV-IM infection may have potential biotechnological applications.

Some previous studies have reported enhanced virulence in *R. solani* strains due to mycovirus infection. The presence of uncharacterized viral dsRNAs is linked to enhanced virulence in *R. solani* (52, 53). Similarly, co-infection with two fungal rhabdoviruses or infection with a fungal mitovirus in *R. solani* AG1-IA strains is associated with increased virulence (54, 55). However, because these fungal strains harbor multiple viruses (at least eight), it remains unclear whether the enhanced virulence results from interactions among mycoviruses in a host with specific genetic backgrounds. Distinctively, our findings demonstrate that RsEV-IM infection is consistently associated with enhanced virulence across diverse *R. solani* strains (AG-3 and AG-1) and plant hosts, irrespective of fungal genetic heterogeneity or co-infection with other mycoviruses. This consistency raises the possibility that RsEV-IM serves as a key virulence determinant in *R. solani* populations. Moreover, RsEV infection enhances *R. solani*’s growth, sclerotia formation, stress tolerance, and dominance over other soil microbes. These improved traits would undoubtedly be advantageous for fungus survival and dissemination. Thus, the comprehensive beneficial effects of RsEV-IM on its host exemplify a well-established mutualistic relationship. Collectively, these observations imply that *R. solani* often establishes mutualistic relationships with various mycoviruses to enhance its biological fitness in nature. These findings are critical for understanding the pathogenicity and ecological traits of *R. solani*, a globally distributed fungal pathogen with significant agricultural and economic impacts.

## Materials and Methods

### Sample collection, isolation of fungal strains and fungal culture

The diseased potato samples were collected from potato fields in the Inner Mongolia Autonomous Region of China from 2008 to 2015. Fungal strains were isolated from the junctions between potato black scurf lesions and healthy tissue using a previously described method (56). For phenotypic growth observation, fungal strains were cultured on potato dextrose agar (PDA) plates at 25°C for 3–5 days. For mycelial collection, strains were grown on PDA medium overlaid with cellophane. The Rs80 strain was cultured on PDA medium for 7 days until sclerotia formation. Individual sclerotia were collected and cultured on PDA medium for 3 days before being subjected to RsEV-IM detection.

For stress-inducing treatments, fresh hyphae were placed on different stress-inducing media for 3–5 days under the same conditions as above. These different stress-inducing media were prepared by adding 10 μg/mL boscalid, 0.1 μg/mL thifluzamide, 1 μg/mL azoxystrobin (fungicides), 1 M NaCl, 0.01% H_2_O_2_, and 1 mM Congo red (final concentration) to PDA medium.

### Plant growth conditions and fungal inoculation

Potatoes and other plants were grown in a greenhouse at 25°C under 16-hour light/8-hour dark cycle conditions. Sterilized toothpicks were used to wound young plant stems. The wounds were covered with 3-day-old fresh mycelial gel plugs (5 mm diameter) with the hyphal side facing inward and wrapped with parafilm. The plate covering the wound was removed after 24 hours. Inoculated plants were maintained in the greenhouse under the same conditions described above, with 80-90% humidity.

For *R. solani* inoculation in soil, first potato seedlings (6 plants) were planted in pots (15 × 45 × 13 cm) containing soil for one week, and then small chopped mycelial PDA plugs fully grown on 10 plates (90 mm diameter) were added to and mixed with the soil.

For mycobiome analysis, soil in pots (15 × 45 × 13 cm) was thoroughly mixed with small chopped mycelial PDA plugs fully grown on 10 plates (90 mm diameter), and then three potato seedlings were planted for three successive growth cycles (total 30 days).

### Total RNA/dsRNA extraction, RT-PCR, and sequencing of viral genome

Total RNA and double-stranded RNA (dsRNA) were isolated as previously described (57). NGS analysis of the dsRNA-enriched fraction was performed on the Illumina HiSeq 4000 platform (Illumina, San Diego, CA, USA) by Hanyu Biotechnology Co., Ltd. (Shanghai, China), following established protocols (58). Bioinformatic analyses were performed as described earlier (59).

For viral genome characterization, first-strand cDNA was synthesized using EasyScript® Reverse Transcriptase (TransGen Biotech, China). The quasi-full-length sequence of RsEV-IM was amplified by RT-PCR with CWBIO DNA polymerase (China) using virus-specific primers designed from NGS-assembled contigs. The 5’- and 3’-terminal sequences of the RsEV-IM genome were verified by RACE (Rapid Amplification of cDNA Ends) (60). All purified PCR products were cloned into the pGEM®-T Easy Vector (Promega, USA) and subjected to Sanger sequencing. The primers used in this study are listed in Table S2.

### RT-qPCR and Northern blot analyses

First-strand cDNA was synthesized as described above. RT-qPCR was performed using PerfectStart Green qPCR SuperMix (Kapa Biosystems, USA) on a CFX96™ Real-Time PCR Detection System (Bio-Rad, USA). Fungal β-tubulin served as internal controls. Three biological replicates were analyzed for each sample, and the experiment was independently repeated three times.

Northern blot analysis was conducted as previously described (61) using digoxigenin (DIG, Roche Diagnostics, Germany)-labeled RNA probes specific to RsEV-IM (RdRp domain).

### Inoculation of RsEV-IM into RsEV-IM-free Fungal Strains

Fresh mycelia from a 3-day PDB culture were finely chopped and placed in a 1.5 mL microcentrifuge tube. They were then incubated with total RNA solution (5 µg/µL) extracted from the Rs80 strain for 30 minutes. After incubation, the mycelia were transferred to fresh PDA medium and cultured. The presence of RsEV-IM in the fungal strains was confirmed by RT-PCR. Strains carrying RsEV-IM were selected and subjected to two successive rounds of similar mycelial RNA treatment to increase virus accumulation levels.

### Isolation of secreted protein fraction and LC-MS/MS analyses

Rs80 and EV-Free strain were cultured in PDB at 25℃ for 3 days. Mycelium were removed by means of miracloth filtration combined with centrifugation at 12,000 g for 20 min, and the liquid supernatant was collected as fungal culture filtrates. Ammonium sulfate was added to the fungal culture filtrates to a final concentration of 40%. After thorough mixing, the mixture was left to stand overnight. Protein precipitation was then obtained by centrifugation at 12,000 rpm for 15 minutes. The protein precipitate was washed with 40% ammonium sulfate solution, then dissolved in 1× PBS.

LC-MS/MS analysis was performed as described previously (62). The raw data files were processed using Proteome Discoverer 2.4 with the Sequest search engine against the *R. solani* proteome (total 12,726 entries). Searches were configured with static modifications for carbamidomethylation of cysteines, dynamic modifications for oxidation of methionine residues and acetylation of protein N-termini, precursor mass tolerance of 10 ppm, fragment mass tolerance of 0.02 Da, and trypsin cleavage (maximum 2 missed cleavages). A reversed sequence decoy strategy was used to control the false discovery rate at ≤ 1% for peptide-spectrum matches, peptides, and proteins.

Label-free quantification was performed using the Minora Feature Detector, Feature Mapper, and Precursor Ions Quantifier nodes in with default settings. Normalization was applied based on the total protein intensity for each sample. Differential protein analysis was performed using the R package *limma*. Putative CWDEs were predicted using dbCAN3 (https://bcb.unl.edu/dbCAN2/blast.php).

### Treatment of fungi/bacteria with siRNA and secreted protein fraction

The siRNA sequences targeting RsEV-IM were synthesized by Tsingke Biological Technology (China), and the assay included not special-siRNA as a negative control. Sequences of siRNAs are provided in Table S2. The siRNA was diluted to 100 nM with 1× PBS buffer (pH 7.4). SiRNAs were used to treat freshly inoculated mycelial plugs placed on PDA medium or mycelial gel plug inoculated to lower stem of plants after one day fungal inoculation.

For treatment using secreted protein, protein concentration was determined using a BCA Protein Quantification Kit-BOX2 from Vazyme (China), and the protein was diluted to 0.25 mg/mL with 1× PBS (pH 7.4) to treat fresh mycelial plugs placed on PDA medium. Overnight-cultured bacteria (100 μL) were pipetted and mixed with an equal volume of the aforementioned secreted protein. After treatment at room temperature for 1 hour, 2 mL of Luria-Bertani (LB) broth was added, followed by shaking culture at 37/28°C for 4 hours. The OD₆₀₀ was measured using an ultraviolet spectrophotometer every hour.

### Eukaryotic protein expression and application to plant

The sequences of three genes, Cellulase, Beta-xylanase, and Pectate lyase, were amplified via RT-PCR. The HA-tagged gene fragments were then cloned into the expression vector pYES2 (digested with *BamH*I and *Hind*III) using the ClonExpress Ultra One Step Cloning Kit V3 from Vazyme (China). Protein expression was performed as previously described (63). The expressed proteins were enriched using Anti-HA Affinity Beads from Smart-Lifesciences (China), dialyzed into 1× PBS (pH 7.4), adjusted to a concentration of 0.1 mg/mL, and injected into the leaves of *Nicotiana benthamiana*. 1× PBS was used as a negative control.

### Sequence and phylogenetic analyses

The Open Reading Frame (ORF) Finder (http://www.ncbi.nlm.nih.gov/gorf/orfg.cgi) program and Conserved Domain Database (CDD, https://www.ncbi.nlm.nih.gov/cdd/) were used for ORF determination and functional unit annotation in proteins. Multiple sequence alignments and result visualization were performed using GeneDoc and ClustalX (version 1.83). Maximum Likelihood (ML) phylogenetic analysis of the RdRP domain was performed using MEGA software (version 10.1.7) with 1000 bootstrap replicates. Conserved motifs in RdRP were predicted by MEME (https://meme-suite.org/meme/tools/meme). *R. solani* proteins were searched at UniProt database (https://www.uniprot.org/).

## Acknowledgements

We thank Ms. Yan Li and Zhimei Bai (Technological Innovation and Talent Cultivation Platform, College of Plant Protection, Northwest A&F University) for technical supports, and Dr. Genhua Yang (Yunnan Agricultural University) for providing research materials. This research was funded by China National Natural Science Foundation (32170163) and Shaanxi Province High-level Talent Grant (F2020225002).

---

**Figure S1.**
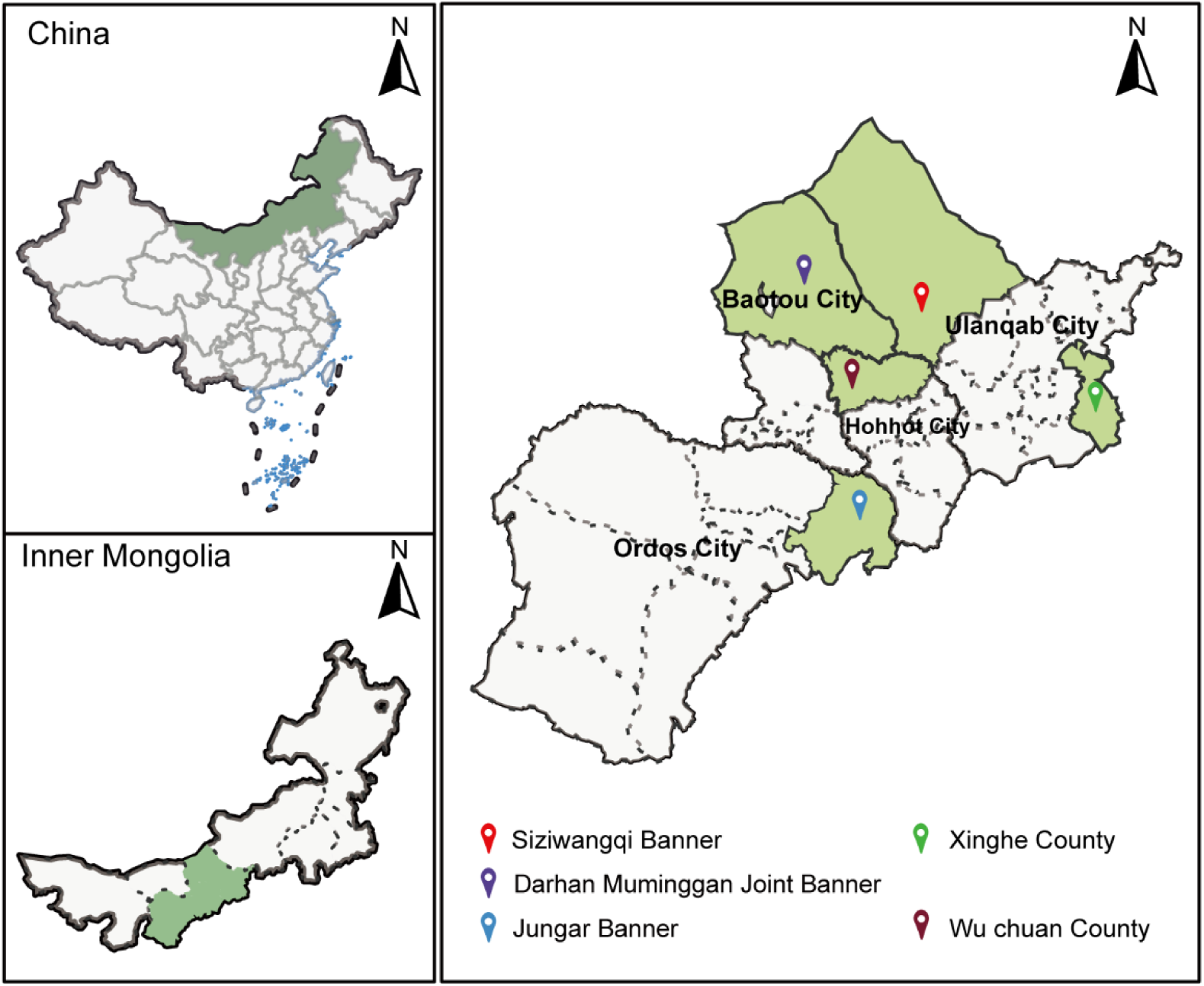
Maps showing the locations where potato samples were collected for the isolation of *R. solani* strains.

**Figure S2.**
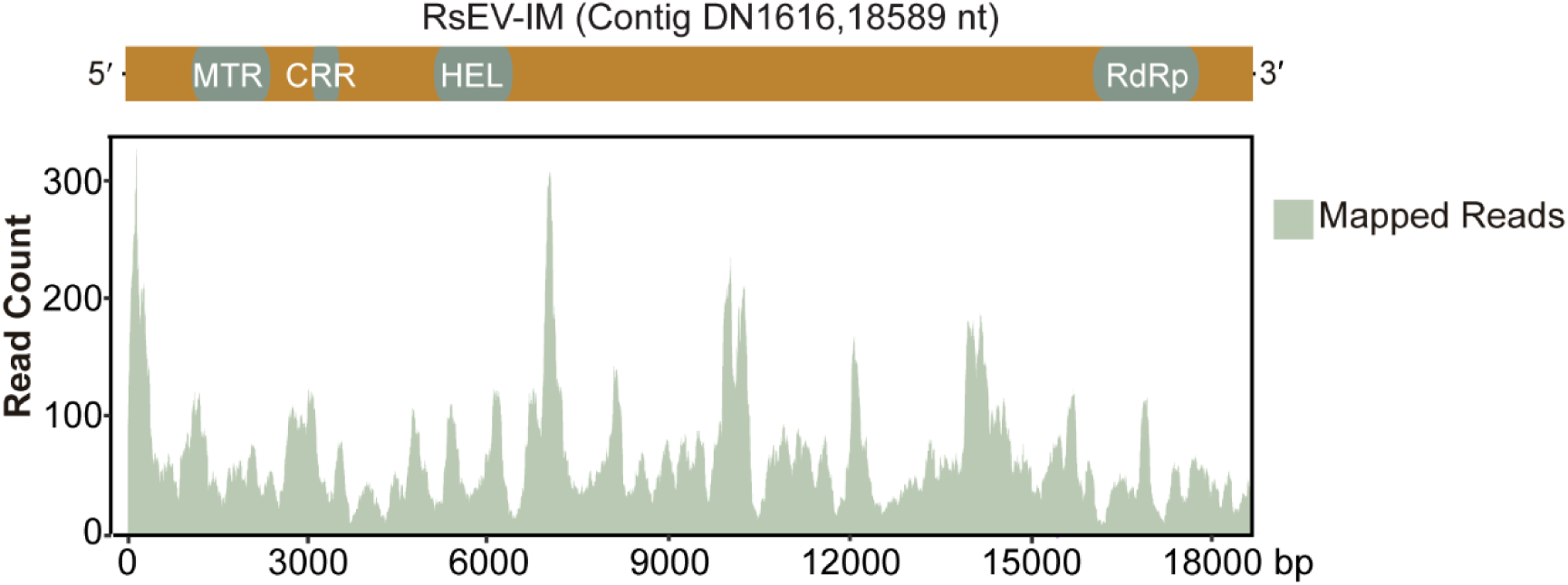
Reads mapped to an endornavirus-related contig.

**Figure S3.**
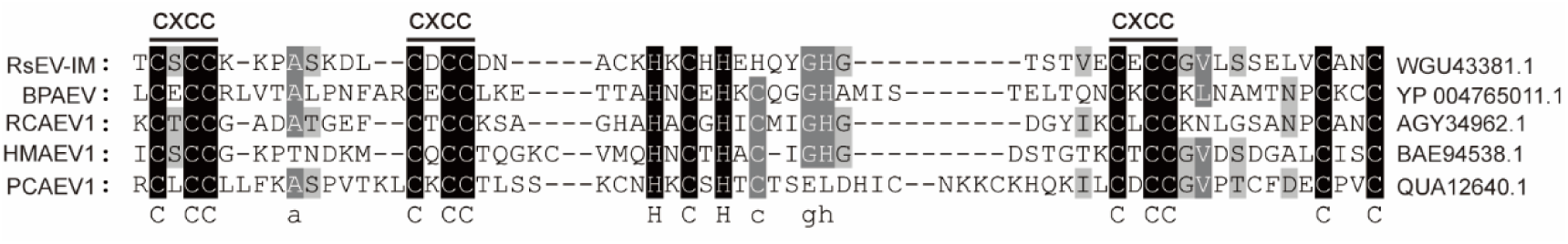
Putative cysteine-rich region (CRR) with conserved "CXCC" signature sequences found in RsEV-IM and other endornaviruses.

**Figure S4.**
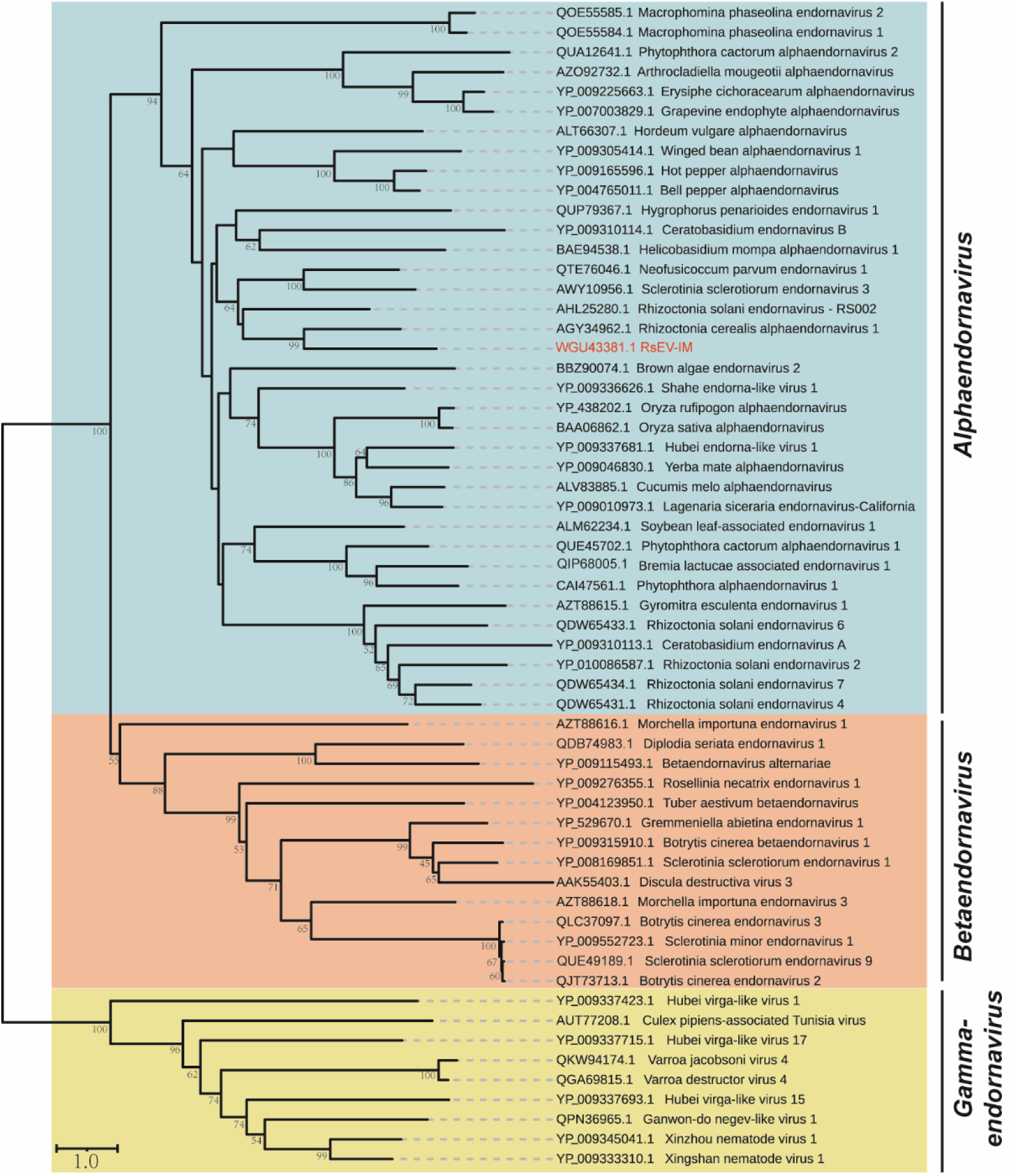
Phylogenetic relationships of RsEV-IM to endornaviruses and selected endorna-like viruses. The tree was constructed using a maximum-likelihood method (JTT matrix-based model) based on a multiple sequence alignment of the RdRp domain. Branch numbers indicate the percentage of replicate trees in which the associated taxa clustered together. The tree is drawn to scale, with branch lengths representing the number of substitutions per site. Virus names are preceded by their accession numbers.

**Figure S5.**
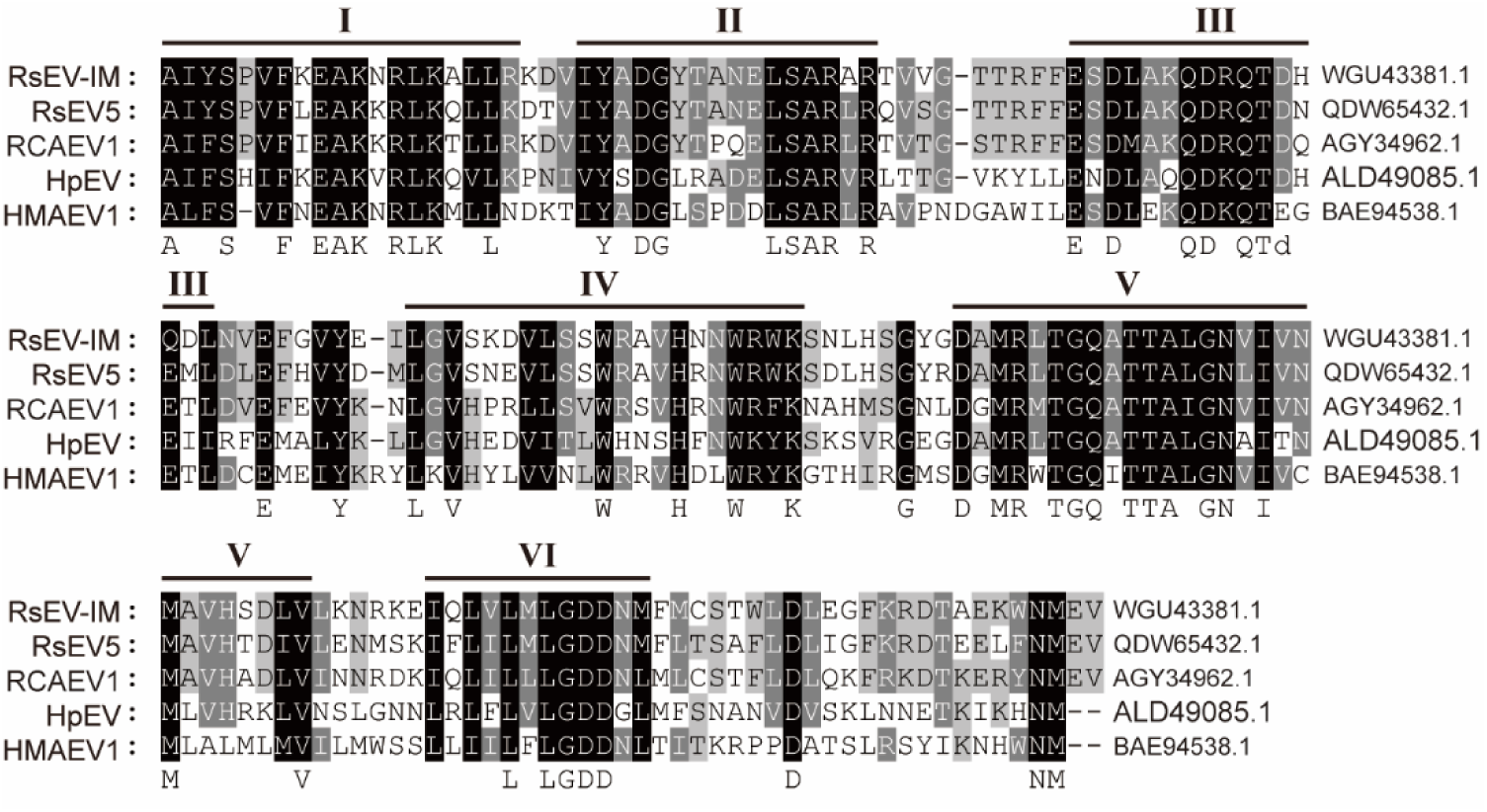
Multiple sequence alignment of the RdRp domain of RsEV-IM with those of alphaendornaviruses.

**Figure S6.**
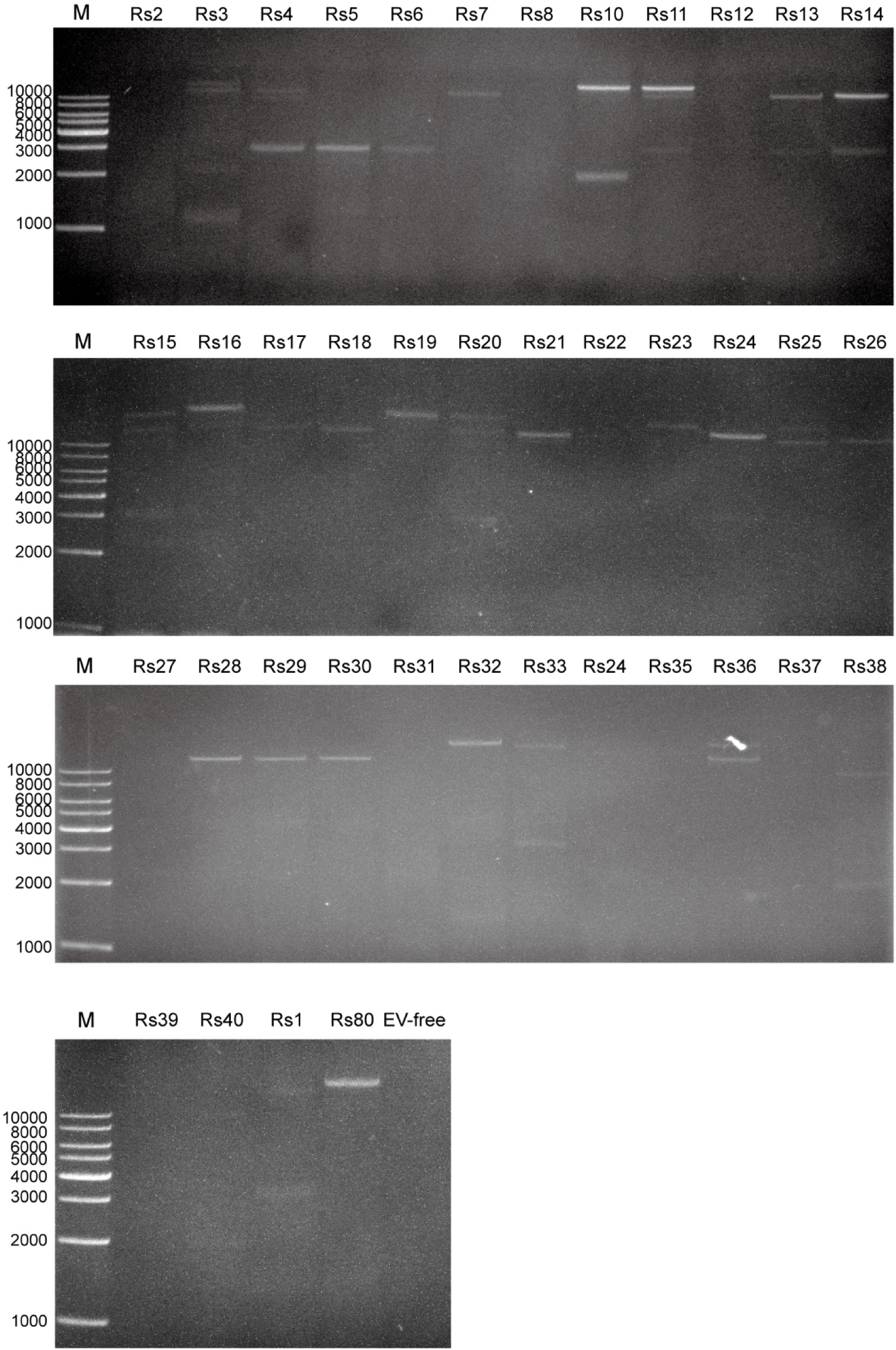
Profiles of dsRNA extracted from *R. solani* AG-3 strains.

**Figure S7.**
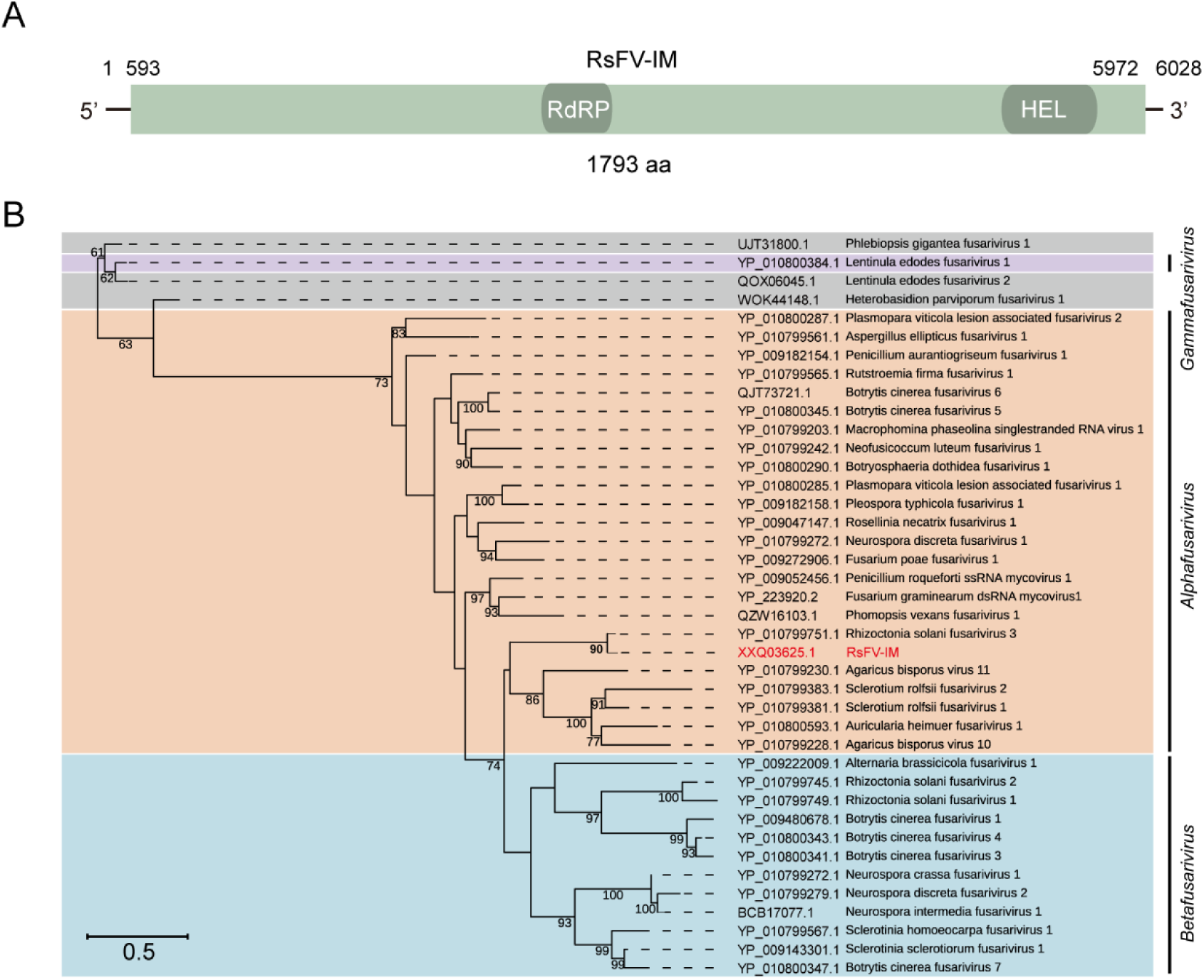
Molecular characteristics and phylogeny of RsFV-IM. **A**. Schematic representation (not to scale) of the RsFV-IM genome structure. Colored boxes represent open reading frames (ORFs), and black lines indicate 5′- and 3′-untranslated regions (UTRs). Nucleotide positions of ORFs and UTRs are labeled. Conserved domains (RdRp and helicase [Hel]) are shown as colored capsule-shaped forms within ORFs. **B**. Phylogenetic relationships of RsFV-IM to other fusariviruses. The tree was constructed using maximum-likelihood (JTT matrix-based model) from an RdRp domain alignment. Branch numbers indicate bootstrap support (%), and branch lengths reflect substitutions per site. Virus names include accession numbers.

**Figure S8.**
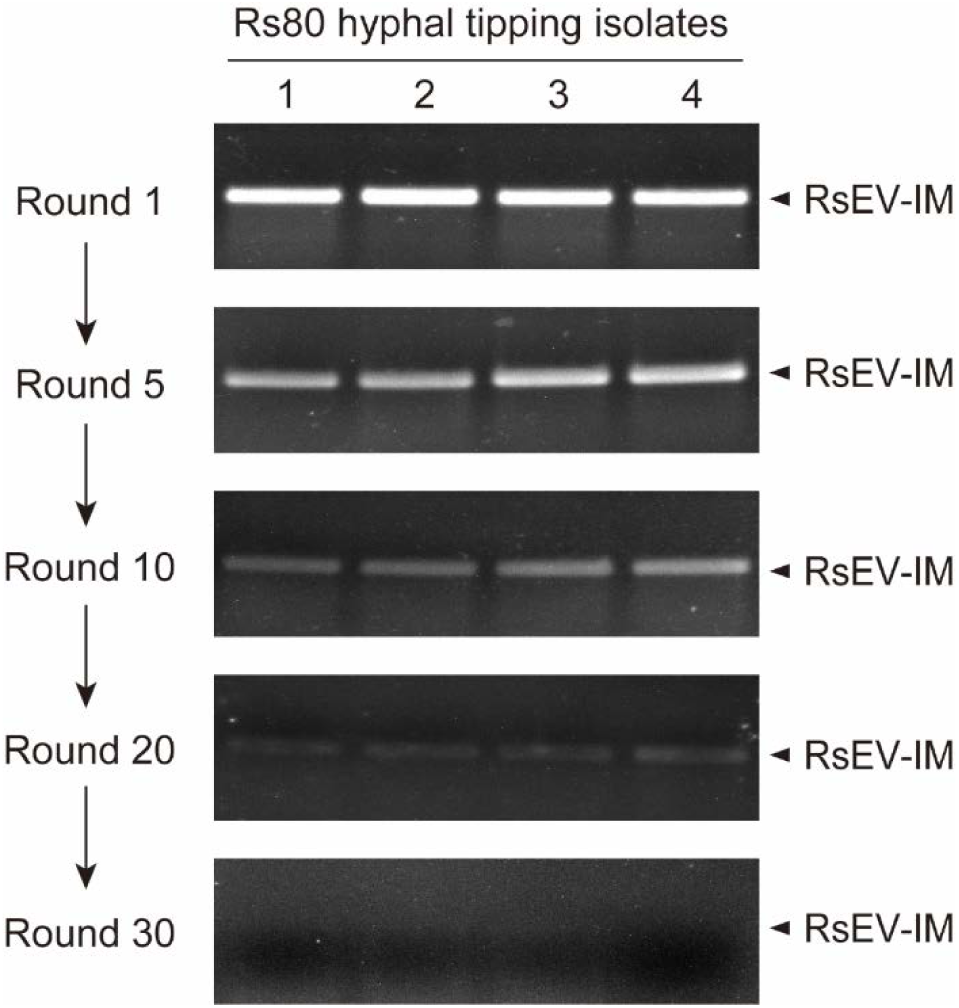
RT-PCR detection of RsEV-IM in fungal isolates derived from successive rounds of hyphal tipping.

**Figure S9.**
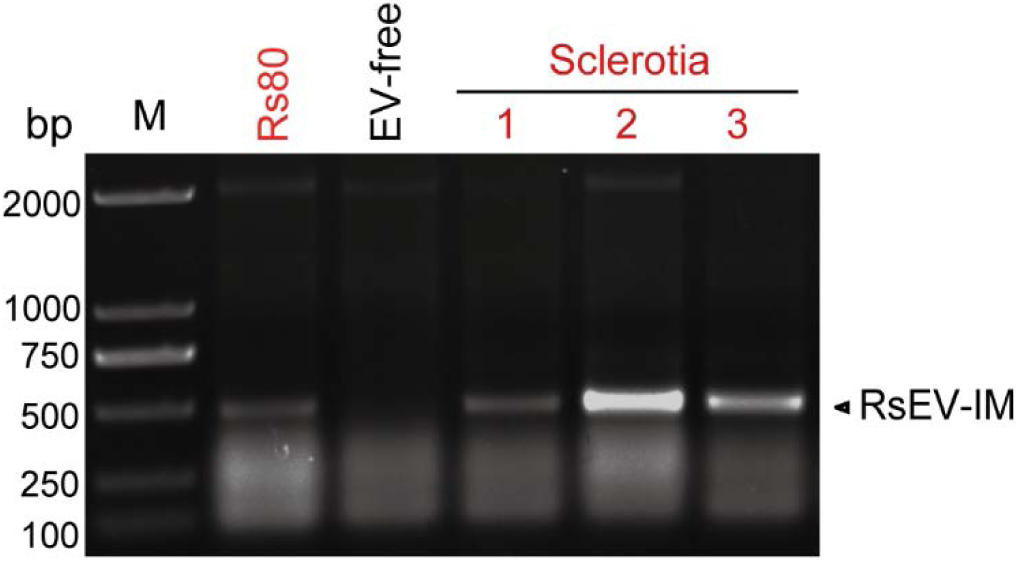
RT-PCR detection of RsEV-IM in fungal isolates regenerated from sclerotia.

**Figure S10.**
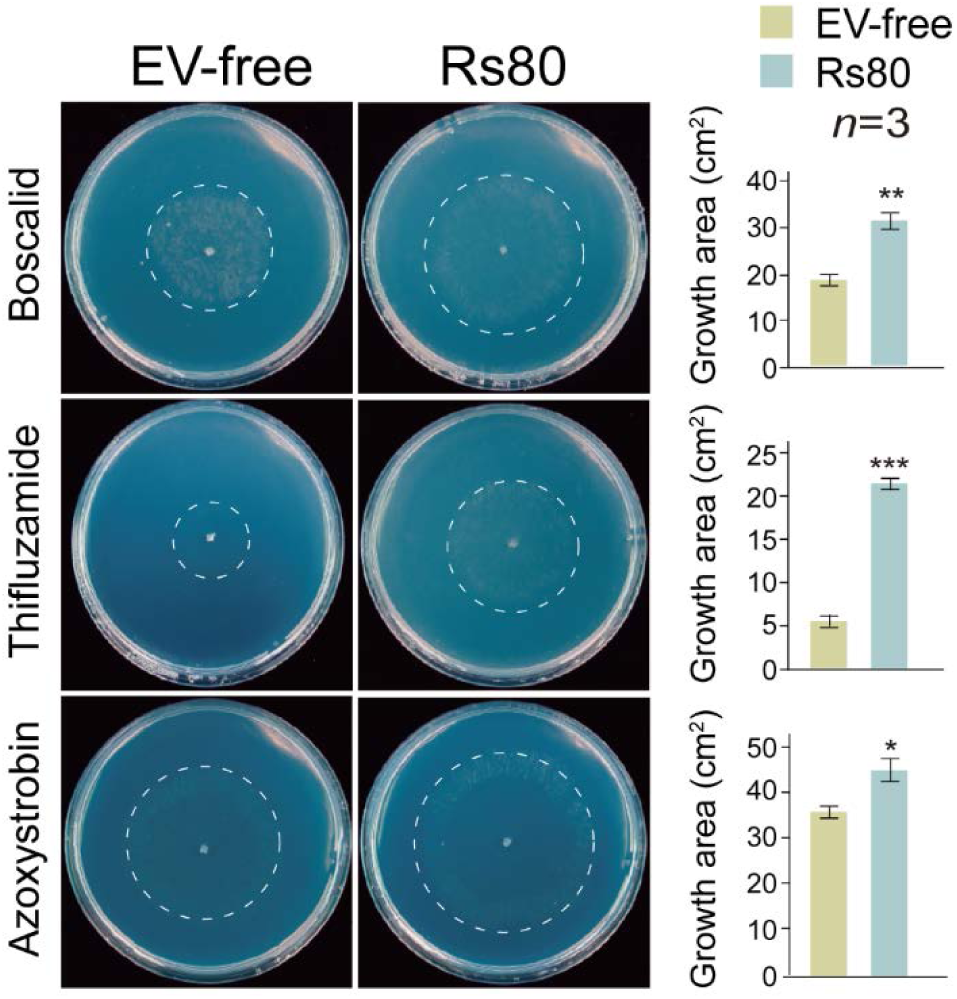
Phenotypic growth of Rs80 and EV-free strains on PDA medium (60 mm plates; imaged at 3 days) supplemented with fungicides. Colony areas (mean ± SD, *n*=3) are shown. Asterisks indicate significance (\**P* < 0.05, \*\**P* < 0.01, \*\*\**P* < 0.001; Student’s t-test).

**Figure S11.**
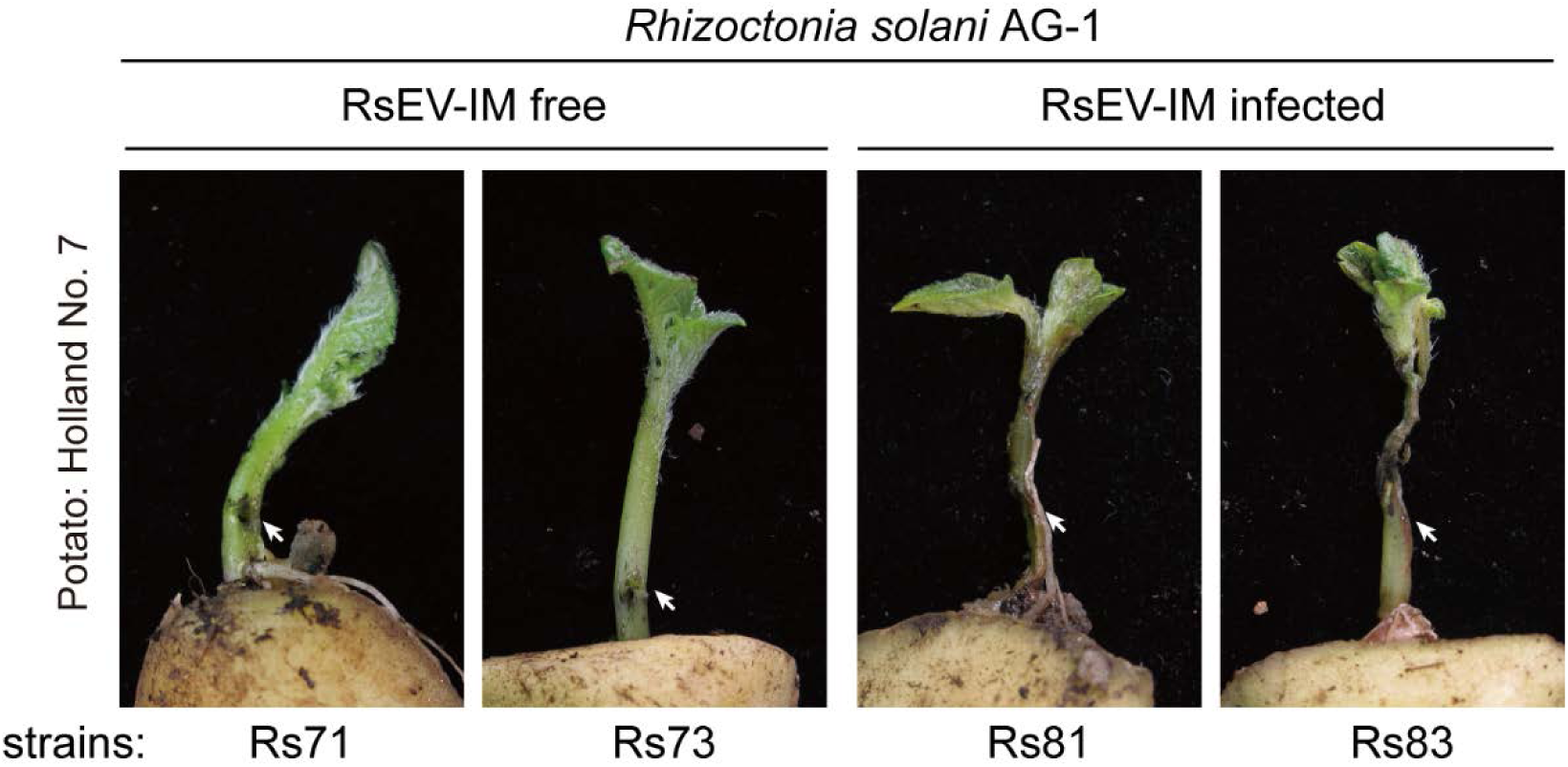
Development of stem rot in lower stems of potato shoots inoculated with RsEV-IM-infected and -free *R. solani* AG-1 strains. Insets present close-up views of the inoculated stems (Magnification: 5×). Plants were photographed at 7 days post-inoculation (dpi). The presented image is representative of three inoculated plants.

**Figure S12.**
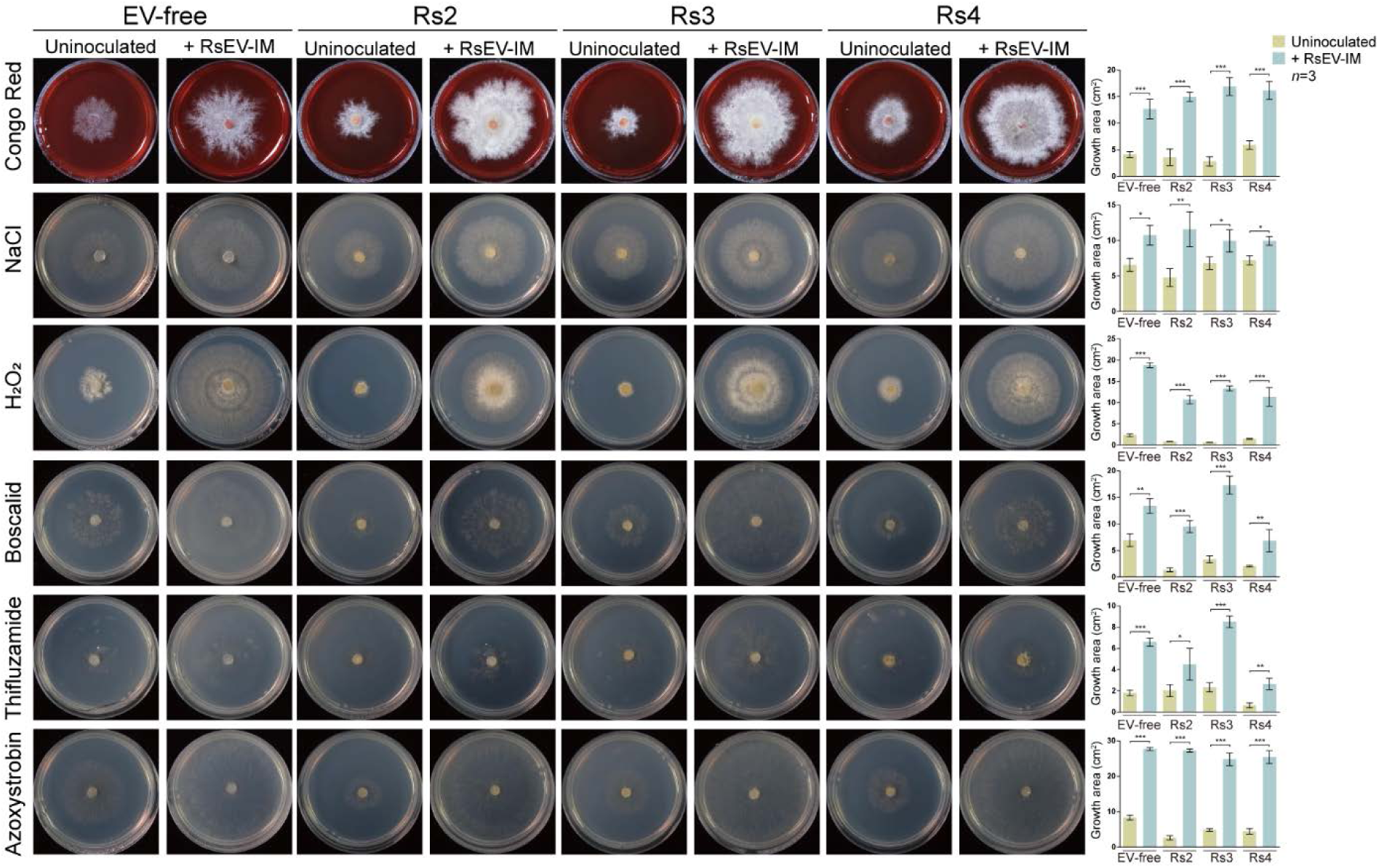
Phenotypic growth of fungal strains that had been inoculated with RsEV-IM on PDA medium (60 mm plates; imaged at 3-4 days) supplemented with stress-inducing agents (congo red; cell wall stress, NaCl; osmotic stress, H2O2; oxidative stress) and fungicides (boscalid, thifluzamide, azoxystrobin). Colony areas (mean ± SD, *n*=3) are shown. Asterisks indicate significance (\**P* < 0.05, \*\**P* < 0.01, \*\*\**P* < 0.001; Student’s t-test).

**Figure S13.**
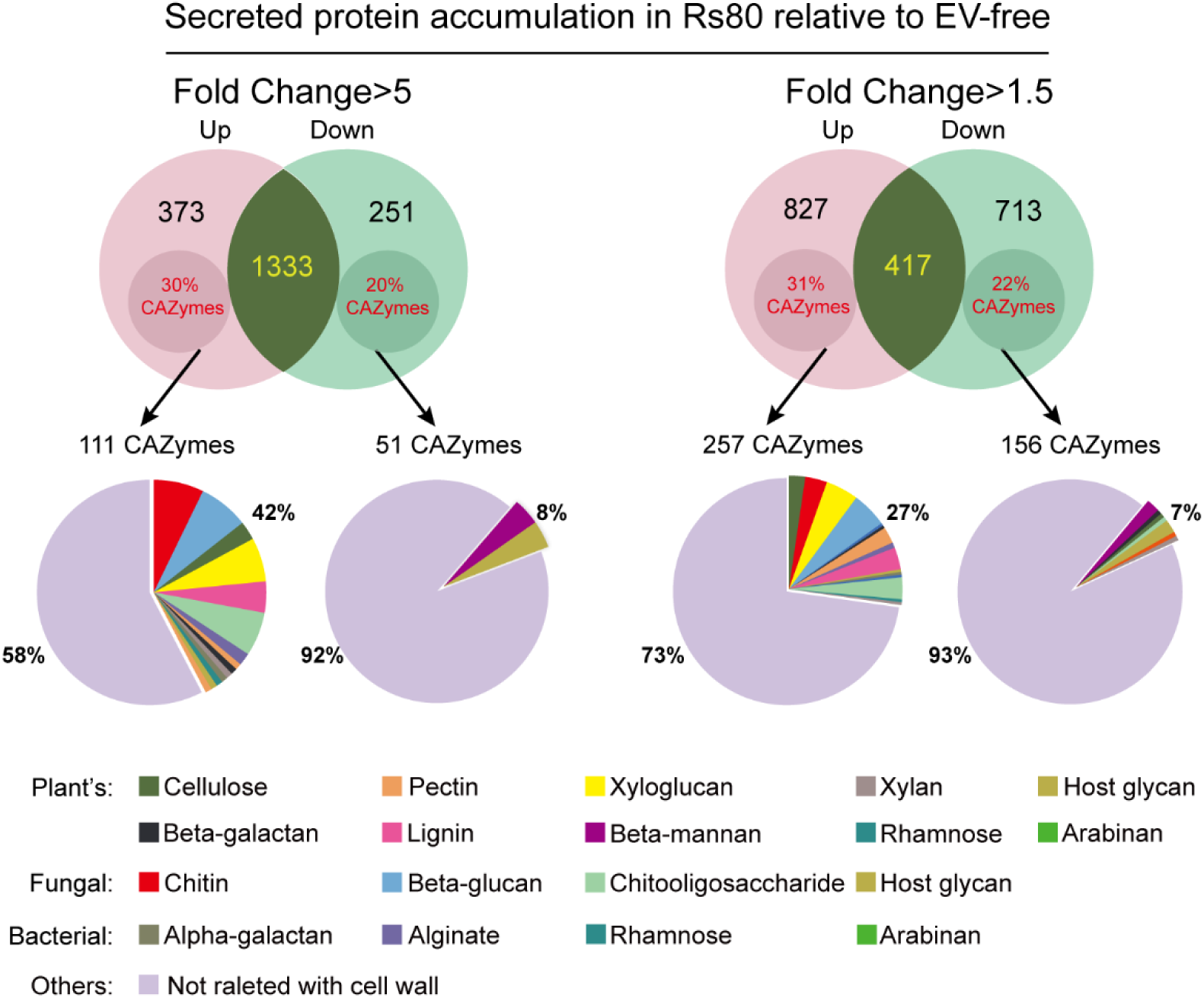
LC-MS/MS analysis of secreted protein fractions from Rs80 and EV-free strains. Proportions of differentially abundant cell wall-digesting enzymes (grouped by substrate specificity) are shown.

**Figure S14.**
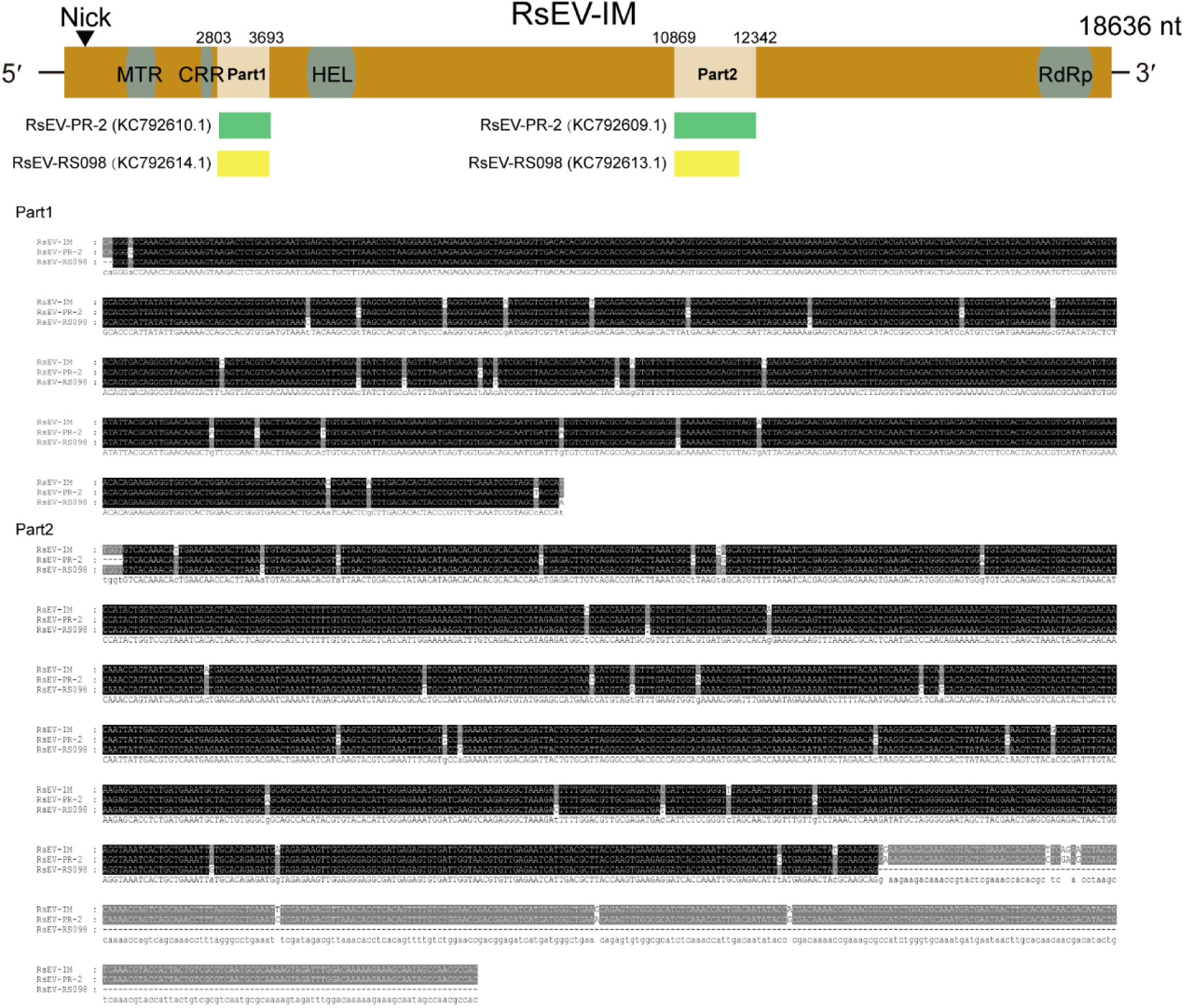
Mapping of partial sequences from dsRNA extracted from *R. solani* AG-3 strains isolated from potato plants in New Zealand to RsEV-IM genome.

**Table S1.**
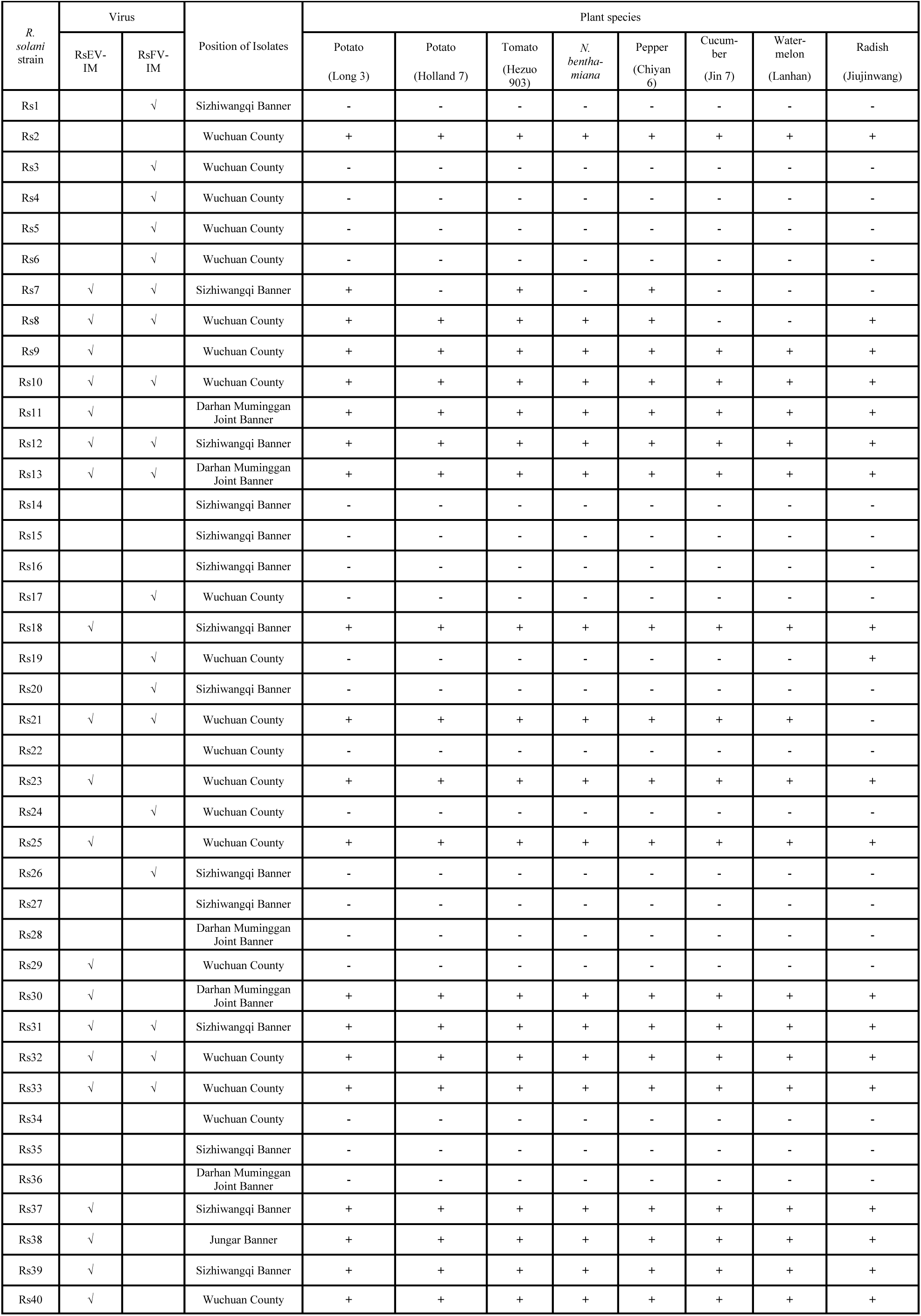
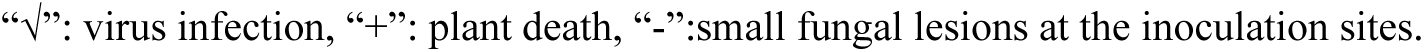
Severe stem rot induced by *Rhizoctonia solani* strains on various plants species.

| *R. solani* strain | Virus |  | Position of Isolates | Plant species |  |  |  |  |  |  |  |
| --- | --- | --- | --- | --- | --- | --- | --- | --- | --- | --- | --- |
|  | RsEV-IM | RsFV-IM |  | Potato<br>(Long 3) | Potato<br>(Holland 7) | Tomato<br>(Hezuo 903) | *N. benthamiana* | Pepper<br>(Chiyan 6) | Cucumber<br>(Jin 7) | Water-melon<br>(Lanhan) | Radish<br>(Jiujinwang) |
| Rs1 |  | √ | Sizhiwangqi Banner | - | - | - | - | - | - | - | - |
| Rs2 |  |  | Wuchuan County | + | + | + | + | + | + | + | + |
| Rs3 |  | √ | Wuchuan County | - | - | - | - | - | - | - | - |
| Rs4 |  | √ | Wuchuan County | - | - | - | - | - | - | - | - |
| Rs5 |  | √ | Wuchuan County | - | - | - | - | - | - | - | - |
| Rs6 |  | √ | Wuchuan County | - | - | - | - | - | - | - | - |
| Rs7 | √ | √ | Sizhiwangqi Banner | + | - | + | - | + | - | - | - |
| Rs8 | √ | √ | Wuchuan County | + | + | + | + | + | - | - | + |
| Rs9 | √ |  | Wuchuan County | + | + | + | + | + | + | + | + |
| Rs10 | √ | √ | Wuchuan County | + | + | + | + | + | + | + | + |
| Rs11 | √ |  | Darhan Muminggan Joint Banner | + | + | + | + | + | + | + | + |
| Rs12 | √ | √ | Sizhiwangqi Banner | + | + | + | + | + | + | + | + |
| Rs13 | √ | √ | Darhan Muminggan Joint Banner | + | + | + | + | + | + | + | + |
| Rs14 |  |  | Sizhiwangqi Banner | - | - | - | - | - | - | - | - |
| Rs15 |  |  | Sizhiwangqi Banner | - | - | - | - | - | - | - | - |
| Rs16 |  |  | Sizhiwangqi Banner | - | - | - | - | - | - | - | - |
| Rs17 |  | √ | Wuchuan County | - | - | - | - | - | - | - | - |
| Rs18 | √ |  | Sizhiwangqi Banner | + | + | + | + | + | + | + | + |
| Rs19 |  | √ | Wuchuan County | - | - | - | - | - | - | - | + |
| Rs20 |  | √ | Sizhiwangqi Banner | - | - | - | - | - | - | - | - |
| Rs21 | √ | √ | Wuchuan County | + | + | + | + | + | + | + | - |
| Rs22 |  |  | Wuchuan County | - | - | - | - | - | - | - | - |
| Rs23 | √ |  | Wuchuan County | + | + | + | + | + | + | + | + |
| Rs24 |  | √ | Wuchuan County | - | - | - | - | - | - | - | - |
| Rs25 | √ |  | Wuchuan County | + | + | + | + | + | + | + | + |
| Rs26 |  | √ | Sizhiwangqi Banner | - | - | - | - | - | - | - | - |
| Rs27 |  |  | Sizhiwangqi Banner | - | - | - | - | - | - | - | - |
| Rs28 |  |  | Darhan Muminggan Joint Banner | - | - | - | - | - | - | - | - |
| Rs29 | √ |  | Wuchuan County | - | - | - | - | - | - | - | - |
| Rs30 | √ |  | Darhan Muminggan Joint Banner | + | + | + | + | + | + | + | + |
| Rs31 | √ | √ | Sizhiwangqi Banner | + | + | + | + | + | + | + | + |
| Rs32 | √ | √ | Wuchuan County | + | + | + | + | + | + | + | + |
| Rs33 | √ | √ | Wuchuan County | + | + | + | + | + | + | + | + |
| Rs34 |  |  | Wuchuan County | - | - | - | - | - | - | - | - |
| Rs35 |  |  | Sizhiwangqi Banner | - | - | - | - | - | - | - | - |
| Rs36 |  |  | Darhan Muminggan Joint Banner | - | - | - | - | - | - | - | - |
| Rs37 | √ |  | Sizhiwangqi Banner | + | + | + | + | + | + | + | + |
| Rs38 | √ |  | Jungar Banner | + | + | + | + | + | + | + | + |
| Rs39 | √ |  | Sizhiwangqi Banner | + | + | + | + | + | + | + | + |
| Rs40 | √ |  | Wuchuan County | + | + | + | + | + | + | + | + |

“√”: virus infection, “+”: plant death, “-”:small fungal lesions at the inoculation sites.

**Table S2.**
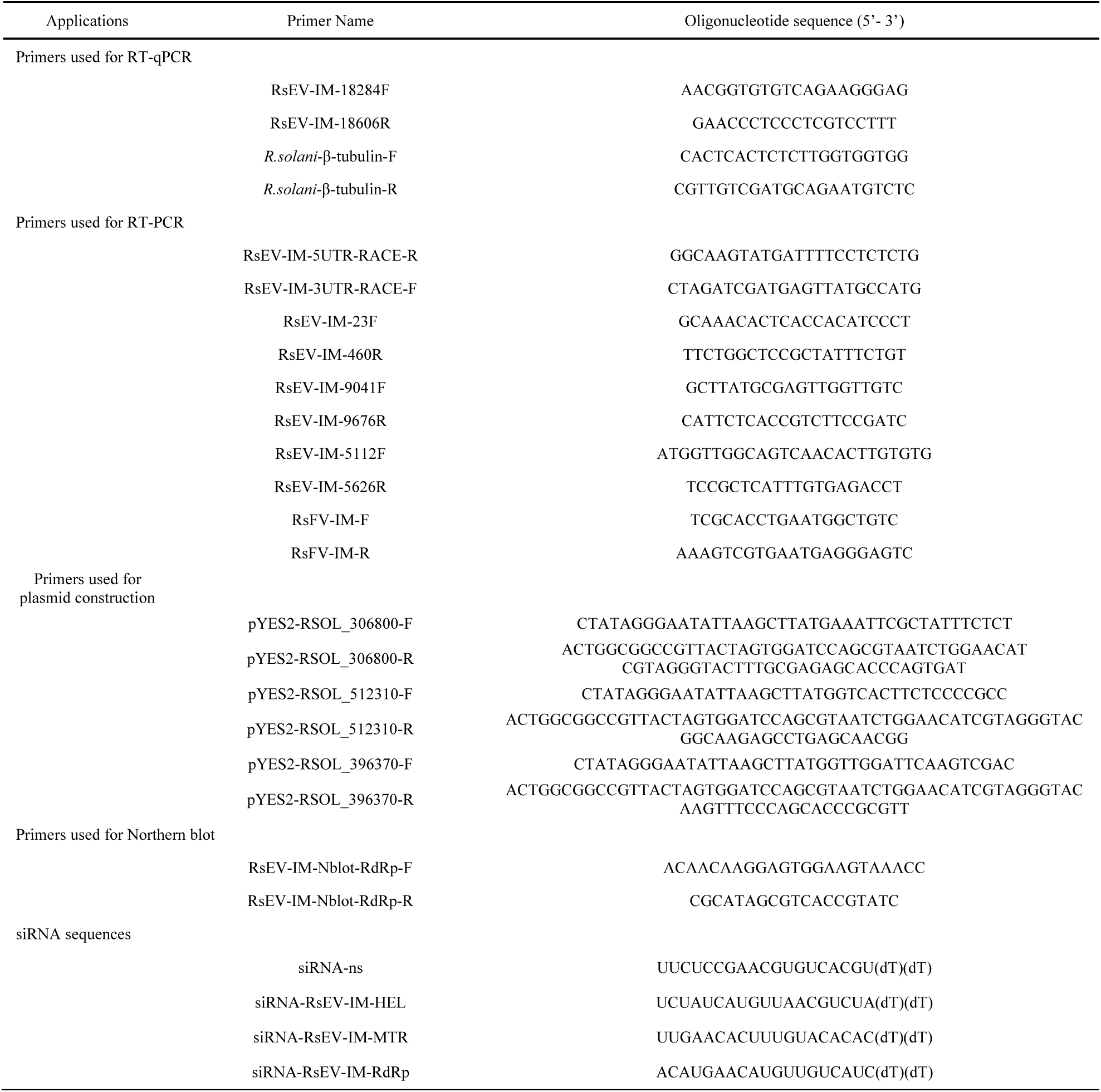
A list of primers and siRNA sequences used in this study.

